# ATR promotes genome instability via CENP-A eviction from centromeres under replication stress

**DOI:** 10.1101/2025.10.20.683416

**Authors:** Denis Ostapenko, Hang Li, Isabelle Trier, Ross Shamby, Eleanor Zagoren, Carla Becerra Sabrera, Kaelyn Sumigray, Lilian Kabeche

**Affiliations:** Department of Molecular Biophysics and Biochemistry Yale University New Haven, CT 06511; Yale Cancer Biology Institute Yale University West Haven, CT 06516; Department of Genetics Yale School of Medicine New Haven, CT 06511

## Abstract

Replication stress leads to genome instability in part by promoting missegregation of chromosomes lacking centromeres. Yet the molecular mechanism linking replication stress to centromere dysfunction has remained elusive. Here, we show that sustained replication stress induces eviction of the histone H3 variant CENP-A. Displaced CENP-A relocalizes to nucleoli. This process is dependent on the DNA damage response kinase, ATR, and occurs in both human and mouse cells. We show that ATR promotes CENP-A eviction by recruiting the AAA+ ATPase VCP to centromeres, destabilizing CENP-A–containing nucleosomes. The canonical CENP-A chaperone, HJURP, but not H3 histone chaperones DAXX or ATRX, is necessary for nucleolar CENP-A localization. Importantly, ATR-dependent CENP-A eviction endures after cell-cycle re-entry and correlates with the emergence of acentric chromosomes, linking replication stress directly to segregation defects. Our findings reveal an undiscovered role for ATR in regulating centromere identity under stress and uncover a mechanistic pathway that drives genome instability.

## INTRODUCTION

Replication stress, arising from stalled or collapsed replication forks that generate single-stranded lesions, leads to elevated chromosome segregation defects ^1–6^. Such defects often manifest as acentric chromosomes or fragments, defined by the absence of centromere and kinetochore proteins. Centromere identity is specified by the histone H3 variant centromere protein A (CENP-A), an essential epigenetic mark that orchestrates kinetochore assembly and thereby ensures faithful chromosome segregation ^7^. Disruption of this mark compromises kinetochore formation and predisposes cells to segregation errors, and the persistence of such errors ultimately drives genome instability^4,8^. While the relationship between replication stress and chromosome missegregation is well established, the mechanism linking these processes remains unknown.

Ataxia telangiectasia and Rad3-related (ATR) kinase, a master regulator of the DNA damage response pathway^9^, safeguards centromere identity^10,11^. Our previous work demonstrated that, in the absence of DNA damage, ATR regulates the association of the histone H3.3 chaperone death domain-associated protein 6 (DAXX) with promyelocytic leukemia nuclear bodies (PML NBs), thereby blocking H3.3 deposition at centromeres^11^. In turn, ATR inhibition releases DAXX and drives the eviction of CENP-A from the centromere. Upon DNA damage, ATR is released from PML bodies^12^. Although DAXX remains at PML bodies after DNA damage (13), its potential role in ATR-dependent CENP-A eviction during replication stress is unknown. Furthermore, in mice, prolonged DNA damage induces CENP-A relocalization from centromeres to nucleoli through an ataxia telangiectasia mutated (ATM)-dependent mechanism that requires the murine-specific CENP-A S30 phosphorylation site^13^. Therefore, it is unclear whether the ATM-dependent pathway has a mechanistic counterpart in humans.

CENP-A deposition and eviction are tightly regulated, with deposition confined to early G1 via Holliday junction recognition protein (HJURP)-mediated incorporation ^14,15^ and eviction triggered under conditions such as DNA damage or replication stress ^6,16,17^. However, how ATR contributes to CENP-A dynamics remains unclear.

Here, we demonstrate that sustained replication stress highly reduces CENP-A occupancy at centromeres, with the displaced CENP-A subsequently relocating to the nucleolus. We further show that this eviction is ATR-dependent but ATM-independent, and that the same pathway operates in mouse cells. Intriguingly, we found that neither DAXX nor its binding partner alpha-thalassemia/mental retardation syndrome X-linked (ATRX) is involved. Instead, HJURP is essential for directing CENP-A to the nucleolus. Finally, we demonstrate that ATR promotes CENP-A eviction via the AAA-ATPase, Vasolin Containing Protein (VCP), and that depletion of ATR rescues the resulting acentric chromosome segregation defects.

## RESULTS

### DNA damage leads to a decrease in CENP-A occupancy

We sought to induce DNA damage using hydroxyurea (HU), an antimetabolite that inhibits ribonucleotide reductase.^18^ 1mM HU treatment for 24 hours led to increased DNA damage, marked by gH2AX and p53 stabilization (**Figure S1A**), suggesting robust DNA damage response activation (**Figure S1A**). A stark decrease in EdU-positive and G2 cells was also observed after treatment with HU (**Figure S1B**). We used chromatin immunoprecipitation coupled with quantitative PCR to measure CENP-A occupancy using primers to the alpha satellite region of three distinct chromosomes, 1, 4, and 15 (**Figure 1A**). HU treatment led to a significant (over 50%) loss of CENP-A occupancy compared to DMSO-treated cells in all chromosomes probed (**Figure 1A**). Because prolonged HU treatment leads to a sustained S-phase arrest, we wanted to make sure that our results were not due to prolonged arrest. Cells were treated with palbociclib, which blocks origin firing and arrests cells at the G1/S entry by inhibiting CDK4 and CDK6 ^19^. In contrast to HU, we did not observe an increase in γH2AX compared to DMSO-treated cells, nor an increase in p53 stabilization (**Figure S1C)**. In contrast to HU treatment, PALB treatment did not significantly decrease the abundance of CENP-A compared to DMSO-treated cells (**Figure 1A**). HU treatment of lung adenocarcinoma A549 cells also led to an increase in p53 and gH2AX (**Figure S1D**). A similar loss of CENP-A occupancy at centromeres occurred after HU treatment using probes for chromosomes 1 and 15 (**Figure 1B**). We sought to confirm these results using additional methods. Cells were fractionated to separate chromatin-bound and unbound (nuclear) pools of CENP-A. A subset (∼20%) of CENP-A was not chromatin-bound in DMSO-treated cells (**Figure 1C, D**). This was similar in PALB- treated cells, where a majority of CENP-A was chromatin-bound (**Figure 1C, D**). In contrast, CENP-A abundance was increased in the nuclear pool (∼53%) in HU-treated cells compared to DMSO-treated cells (**Figure 1C, D**). These data are consistent with work in non-cancer cell lines, RPE-1 cells^6,17^. Importantly, we did not observe an aberrant decrease in CENP-A total abundance after HU treatment, suggesting that CENP-A loss at centromeres was not leading to its aberrant degradation under these conditions (**Figure 1E**). Taken together, these data demonstrate that CENP-A occupancy at centromeres is decreased after sustained DNA damage.

**Figure 1:**
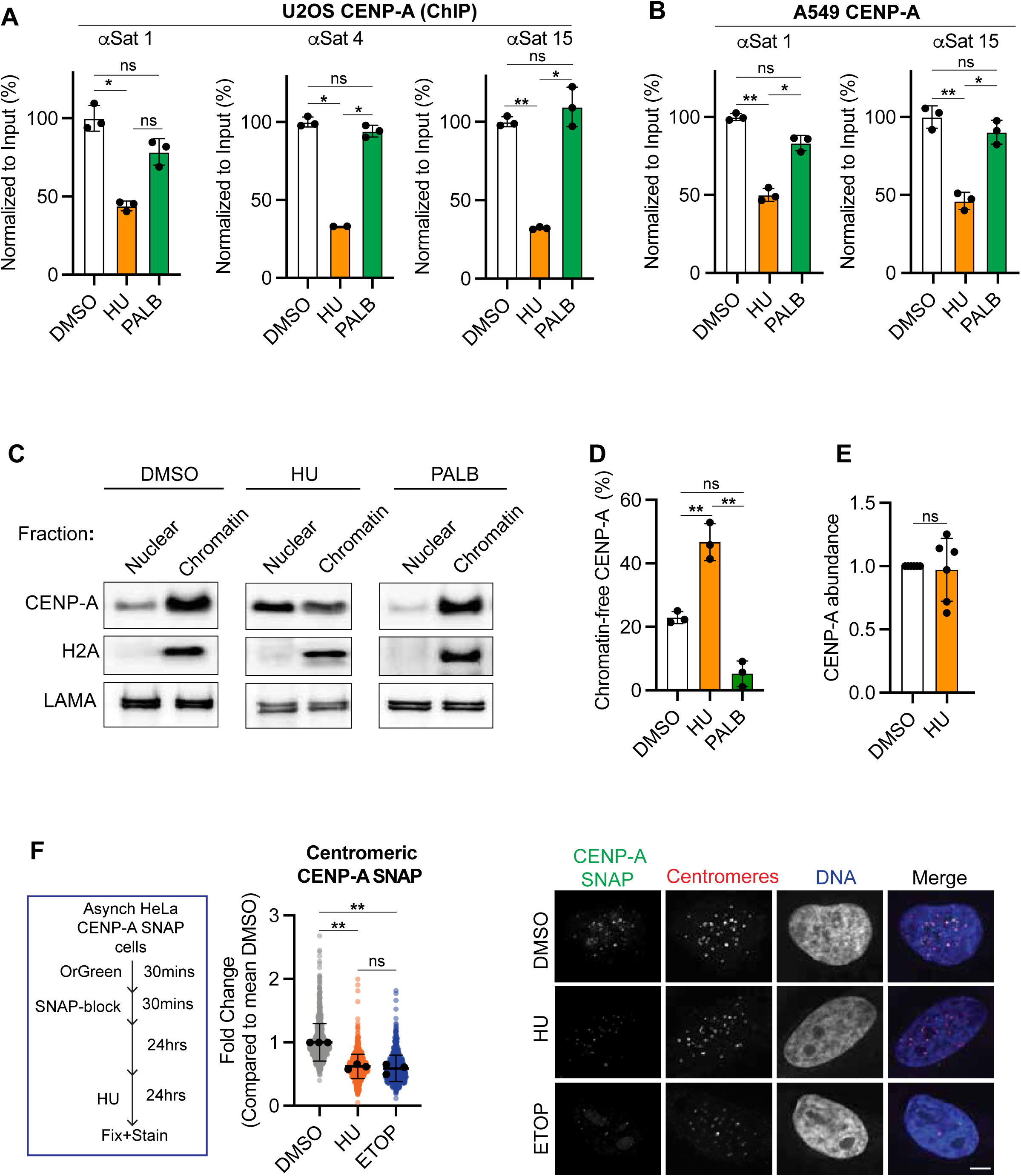
DNA damage leads to a decrease in CENP-A occupancy: (**A**) CENP-A occupancy at the centromeric regions corresponding to chromosomes 1 (αSat1), 4 (αSat4), and 15 (αSat15). Quantitative PCR of CENP-A chromatin immunoprecipitation from U2OS cells treated for 24 hr with either DMSO, 1 mM hydroxyurea (HU), or 0.5 µM palbociclib (PALB). Black points represent independent replicates with mean and SD. (**B**) CENP-A chromatin immunoprecipitation from A549 cells after 24 hr treatment with DMSO, 1 mM HU, or 0.5 µM PALB. qPCR was carried out with primers corresponding to chromosomes 1 (αSat1) and 15 (αSat15). (**C**) Nuclear and chromatin-containing fractions of U2OS cells pre-treated for 24 hr with DMSO, 1 mM HU, or 0.5 µM PALB were separated on SDS-PAGE and blotted for the presence of CENP-A (upper panel), histone H2A (middle panel), and Lamin A/C (lower panel). (**D**) The intensity of CENP-A Western blots from individual fractions of U2OS cells in (C) was quantified, normalized, and plotted to reflect the chromatin-free CENP-A pool. Black points represent independent replicate averages. (**E**) The total intensity of CENP-A protein blots from U2OS cells treated with DMSO or 1 mM HU for 24 hr was quantified and plotted. (**F**) Schema and quantification (left) and representative images (right) of the intensity of “ancestral” CENP-A SNAP at centromeres in stably expressing HeLa cells after treatment with DMSO, 1 mM HU, or 3 µM etoposide ETOP. SNAP-CENP-A was labeled 24 hr prior to drug treatment. Centromeres were marked using ACA. Scale bar = 5 µm. Black points represent the mean replicate averages, and error bars represent SD. *p < 0.05; **p < 0.01. Significance was calculated using the Student’s t-test where there were two conditions only (E), or one-way ANOVA with Dunnett’s test when comparing multiple conditions to DMSO (A–D, F).

CENP-A deposition is tightly regulated, with deposition occurring in early G1 (ref). Because cells were treated for 24 hours, we could not rule out the possibility that the observed decrease in CENP-A occupancy was due to CENP-A not being properly deposited in G1 before the arrest.

We used CENP-A SNAP-expressing HeLa cells to test this possibility, where CENP-A was stably expressed at near endogenous levels ^20^. Similar to U2OS cells, HeLa cells treated with hydroxyurea (HU) had increased γH2AX abundance compared to DMSO-treated cells(**Figure S1E**). We also treated cells with etoposide, a topoisomerase II inhibitor,^21^ which caused a similar increase in DNA damage (marked by γH2AX) and arrest compared to HU (**Figure S1A-B,1E**) Figure S1A-B; S1E). CENP-A SNAP HeLa cells were incubated with OrGreen for 30 minutes, 24 hours prior to the addition of HU, ETOP, or DMSO. HU and ETOP-treated cells showed an overall decrease (∼40%) in OrGreen-marked CENP-A SNAP at centromeres in ETOP-treated cells compared to DMSO-treated cells (**Figure 1F**). These data suggest that prolonged DNA damage leads to decreased CENP-A abundance at centromeres.

The loss of CENP-A at centromeres led us to assess if CENP-A occupancy was due to general DNA damage at centromeres, causing complete loss of nucleosomes due to prolonged DNA damage, as has been observed in other cell lines^6^. While there was a stark decrease in CENP-A occupancy at centromeres in HU-treated cells that was not present in PALB-treated cells (**Figure 1A**), H2A after HU treatment remained at similar levels to PALB-treated cells at the three centromere regions we tested (**Figure S1G**), suggesting that loss of whole nucleosomes cannot completely account for the loss of CENP-A occupancy. Consistent with these results, there was a moderate but non-significant increase in Rad51 at centromeres compared to DMSO-treated cells in HU and ETOP-treated cells (**Figure S1F**). Overall, these data demonstrate that prolonged DNA damage leads to specific CENP-A eviction from centromeres.

### CENP-A relocalizes to nucleoli after prolonged DNA damage

In addition to decreased CENP-A occupancy at centromeres, we also observed diffuse localization of CENP-A in the nucleus, which overlapped with the nucleolar protein UBTF, an RNA Polymerase I transcription factor, and a component of the nucleolus^22^, after HU or ETOP treatment (**Figure 2A, Figure S2A**). In contrast, this was not observed in palbociclib-treated cells (**Figure S2B**), suggesting that CENP-A localization to nucleoli occurred only after persistent DNA damage. We sought to quantify the percentage of cells that were positive for nucleolar CENP-A. To do this, we developed a macro to isolate individual nuclei and calculate a Pearson’s correlation coefficient (R) between nuclear CENP-A and UBTF immunofluorescence intensity (see methods). In HU or ETOP-treated cells, we observed a significant shift in Rob values compared to DMSO-treated cells (DMSO = 0.43; HU = 0.59; ETOP = 0.57), across multiple biological replicates (**Figure S2C**). In contrast, observed similar Rob values for palbociclib-treated cells (PALB) with a mean of 0.18 across three biological replicates (**Figure S2D**). We decided to use a Rob value of 0.65 to delineate positive co-localization between CENP-A and UBTF, as it exceeded the standard deviation observed in DMSO and PALB-treated cells. This was also where we could concretely observe nucleolar CENP-A localization. Using this metric, we calculated that approximately 11% of DMSO-treated cells were positive for CENP-A localization to the nucleolus (**Figure 2B**). This was significantly increased in HU and ETOP-treated cells, with approximately 38% and 40% of cells, respectively, exhibiting this nucleolar CENP-A phenotype (**Figure 2B**). In contrast, PALB treatment did not lead to a significant change in the percentage of cells with nucleolar CENP-A compared to DMSO-treated cells (**Figure 2C**). Moreover, we observed this nucleolar CENP-A phenotype using an additional monoclonal mouse antibody against the N-terminus of CENP-A (**Figure S2E**), demonstrating that this was not an artifact of the CENP-A antibody.

**Figure 2.**
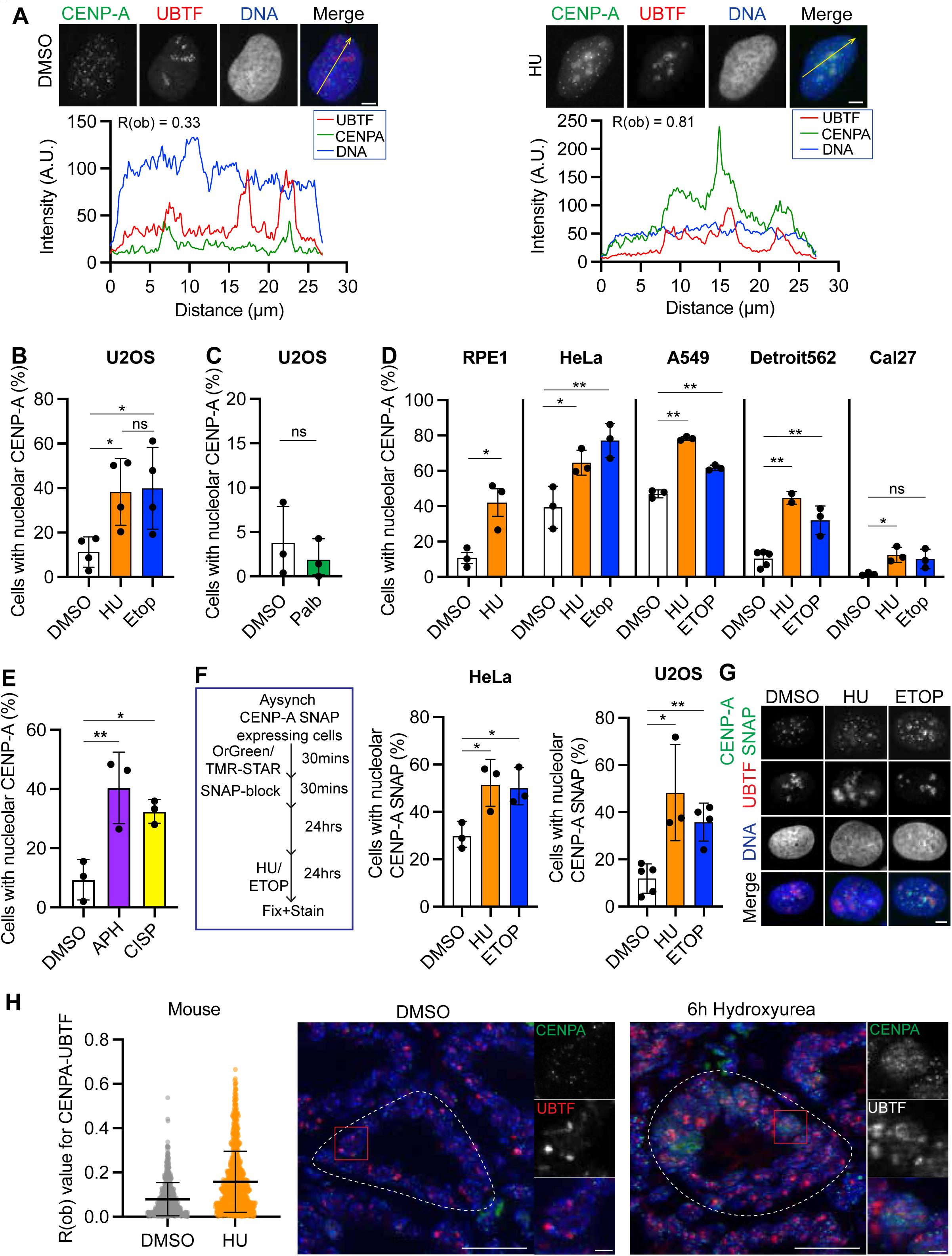
CENP-A relocalizes to nucleoli after prolonged DNA damage: (**A**) Representative image (top) and raw pixel intensity of CENP-A (green), UBTF (red) and DNA (blue) of U2OS cell that was treated with either DMSO (left) or 1mM Hydroxyurea(HU) (right) for 24 hours. R(ob) is also shown for each cell. (**B**) Percentage of U2OS cells that exhibit CENP-A localization to the nucleolus (marked by UBTF) after 24-hour treatment with either DMSO, 1mM Hydroxyurea (HU), or 3μM Etoposide (ETOP). (**C**) Percentage of U2OS cells that exhibit CENP-A localization to the nucleolus (marked by UBTF) after 24-hour treatment with either DMSO or 1μM Palbociclib (PALB). (D) Percentage of cells that exhibit CENP-A localization to the nucleolus (marked by UBTF) after 24-hour treatment with either DMSO, HU, or ETOP. (**E**) Percentage of cells that exhibit CENP-A localization to the nucleolus (marked by UBTF) after treatment with either DMSO, 1μM Aphidicolin (APH), or 1μM Cisplatin (CISP). (**F**) Schema (left) and quantification (right) of the percent of cells that were positive for CENP-A SNAP localization to the nucleolus (marked by UBTF). CENP-A SNAP was labeled prior to treatment. (**G**) Representative image of CENP-A SNAP expressing U2OS cells that were treated with either DMSO or HU whereby CENP-A SNAP was labeled prior to treatment (green), and stained for UBTF (red), and DNA (blue). Scale bar = 5μm. (**H**) Quantification of R(ob) values (left) and representative images (right) of mouse small intestinal crypt cells after treatment with DMSO or 50 mg/kg Hydroxyurea (HU) for 6 hours. Tissues were stained for CENP-A (green), UBTF (red) and DNA (blue). Scale bar = 20 μm on large image and scale bar on inset of single cell = 5μm. Black points represent the mean replicate averages, and error bars represent standard deviation. *p<0.5; **p<0.01. Significance was calculated using the Student-T test, where there are two conditions only (C, H, RPE1 in D), or ANOVA with Dunnett test when comparing multiple conditions to DMSO (D, E, F), or Tukey test when comparing across all conditions (B).

To confirm the specificity of CENP-A localization to nucleoli, we inhibited RNA Pol I using BMH-21. 24-hour treatment of 1 μM BMH-21 led to the visual disintegration of nucleoli (**Figure S2F**). Under these conditions, neither HU nor ETOP treatment led to a considerable increase in this phenotype (**Figure S2E**). These data support our observation that CENP-A localizes specifically to nucleoli. Taken together, these data demonstrate that long-term HU or ETOP treatment leads to increased CENP-A localization to nucleoli.

We sought to determine if this effect was specific to U2OS cells. We extended this analysis to RPE1 (non-cancer retinal epithelial cell line) and several cancer cell lines: HeLa (cervical cancer, p53-, ATL-, HPV+), A549 (lung adenocarcinoma, p53+, HPV-), Detroit 562 (pharyngeal carcinoma,p53+, HPV-), Cal 27 (squamous cell carcinoma,p53+, HPV-). Cell lines had varying basal levels of percentage of cells exhibiting nucleolar CENP-A positive, with Cal27 having the lowest percentage (∼2.5%) to A549 with the highest (∼47%) (**Figure 2D**). All cell lines exhibited a significant increase in the percentage of cells positive for nucleolar CENP-A when treated with either HU or ETOP for 24 hours (**Figure 2D**). We further sought to determine if incubation of U2OS cells with a variety of chemotherapies that have been shown to induce DNA damage, including aphidicolin, cisplatin, and X-ray irradiation (ref). U2OS cells were treated with either aphidicolin (Concentration) or Cisplatin (Concentration) for 24 hours, or irradiated and fixed 24 hours after. We observed that all three treatments led to a significant increase in the percentage of cells that were positive for nucleolar CENP-A (**Figure 2E, Figure S2G)**. Consistent with our findings, data using proximity labeling also found increased interactions with nuclear proteins after replication stress induction in RPE-1 cells using aphidicolin ^17^. Taken together, these data demonstrate that prolonged DNA damage treatment leads to increased CENP-A localization to the nucleolus.

Human CENP-A synthesis is cell-cycle-regulated, with a peak of synthesis in G2^20^. Newly synthesized CENP-A has been observed to be similar to and localized to the nucleolus and interacts with nuclear protein NPM1 ^15,20^. This left the possibility that the pool of nucleolar CENP-A observed after replication stress was newly synthesized CENP-A and not from the displaced CENP-A. To address this, we used U2OS and HeLa CENP-A SNAP-expressing cells that were incubated with OrGreen or TMR-STAR, as in Fig. 1F, and measured the abundance of CENP-A-SNAP at nucleoli using UBTF as a marker for nucleoli. Under these conditions, we observed a strong signal of labeled CENP-A SNAP in HeLa and U2OS cells, which colocalized with UBTF after HU or ETOP treatment (**Figure 2F**). Using correlation analysis, we observed a strong increase in the percentage of cells that were positive for ancestral CENP-A in the nucleolus (**Figure 2F**). We observed that when nascent CENP-A was labeled by pulsing SNAP label after DMSO or HU treatment just before fixing cells, many cells were positive for nucleolar CENP-A SNAP. Still, there was no significant difference between DMSO, HU, or ETOP-treated cells (∼46%, 68%, 49%, respectively) (**Figure S2H**). Taken together, these data demonstrate that ancestral CENP-A, which was previously localized to centromeres, is relocalized to nucleoli after long-term DNA damage.

While we observed strong changes in CENP-A localization in cellulo after treatment with HU or ETOP, we sought to determine if we could also observe CENP-A localization to nucleoli in vivo. We treated a total of 6 mice (postnatal day 20) from two litters of mice with 50 mg/kg of hydroxyurea (HU) for 6 hours and subsequently isolated their small intestines. Tissues were stained for CENP-A, UBTF, and DNA, and R(ob) was measured on nucleoli located at small intestinal crypts, as those represent cycling cells. There was a significant increase in correlation between CENP-A and UBTF in DMSO-treated mice, as measured by R(ob) in HU-treated mice (mean of 0.15), compared to DMSO-treated mice (mean of 0.05) (**Figure 2H**). These data suggest that replication stress leads to an increase in CENP-A localization to nucleoli *in cellulo* and *in vivo*.

### CENP-A relocalization to nucleoli relies on ATR activity

The DNA damage response relies on the activation of master kinases, including ATM and ATR. Their activity is highly dependent on the type of lesion and upstream recognizers of DNA damage^23^. Etoposide and Hydroxyurea can induce DNA damage and cause replication stress and single-stranded DNA lesions, and are often characterized by activation of ATR kinase activity^1^. However, a similar phenotype of CENP-A relocalization in response to DNA damage was observed in mouse cells after prolonged treatment with ETOP. Under these conditions, ETOP led to the relocalization of CENP-A in an ATM dependent ^24^manner. Delocalization was dependent on CENP-A S30, a putative ATM phosphorylation site, which is not present in human cells^24^. Thus, we sought to resolve whether CENP-A eviction and subsequent relocalization to nucleoli occurred due to a specific type of DNA damage or DNA damage response, and if this differed between humans and mice. We titrated etoposide (ETOP), ranging from 80nM to 80 μM, for 24 hours and measured DNA damage using γH2AX. Importantly, γH2AX is a strong marker for ATM activity and double-stranded breaks, as γH2AX is predominantly phosphorylated by ATM^25^. There was a small increase in the γH2AX abundance when cells were treated with 80nM and 0.8 μM ETOP (∼1.3 and 3.3 fold, respectively), a more drastic (∼25 fold) increase when treated with 8 μM ETOP (**Figure 3A, Figure S3A**). There was an even more drastic (∼96-fold) increase after 80μM ETOP treatment (**Figure 3A, Figure S3A**). We also measured replication stress under these conditions by identifying S-phase cells using DAPI and then measuring the intensity of EdU in those cells (**Figure S3B**). Unlike γH2AX abundance, the replication stress score was highest in 0.8 μM ETOP-treated cells and was not significantly increased in 80μM ETOP-treated cells compared to DMSO cells (**Figure 3B**). Similarly, 0.8μM exhibited the highest percentage of cells with nucleolar CENP-A, while 80 μM showed similar rates to DMSO-treated cells (**Figure 3C, Figure S3C**). These data suggest that relocalization of CENP-A to nucleoli occurs due to replication stress and not due to the abundance of DNA damage.

**Figure 3:**
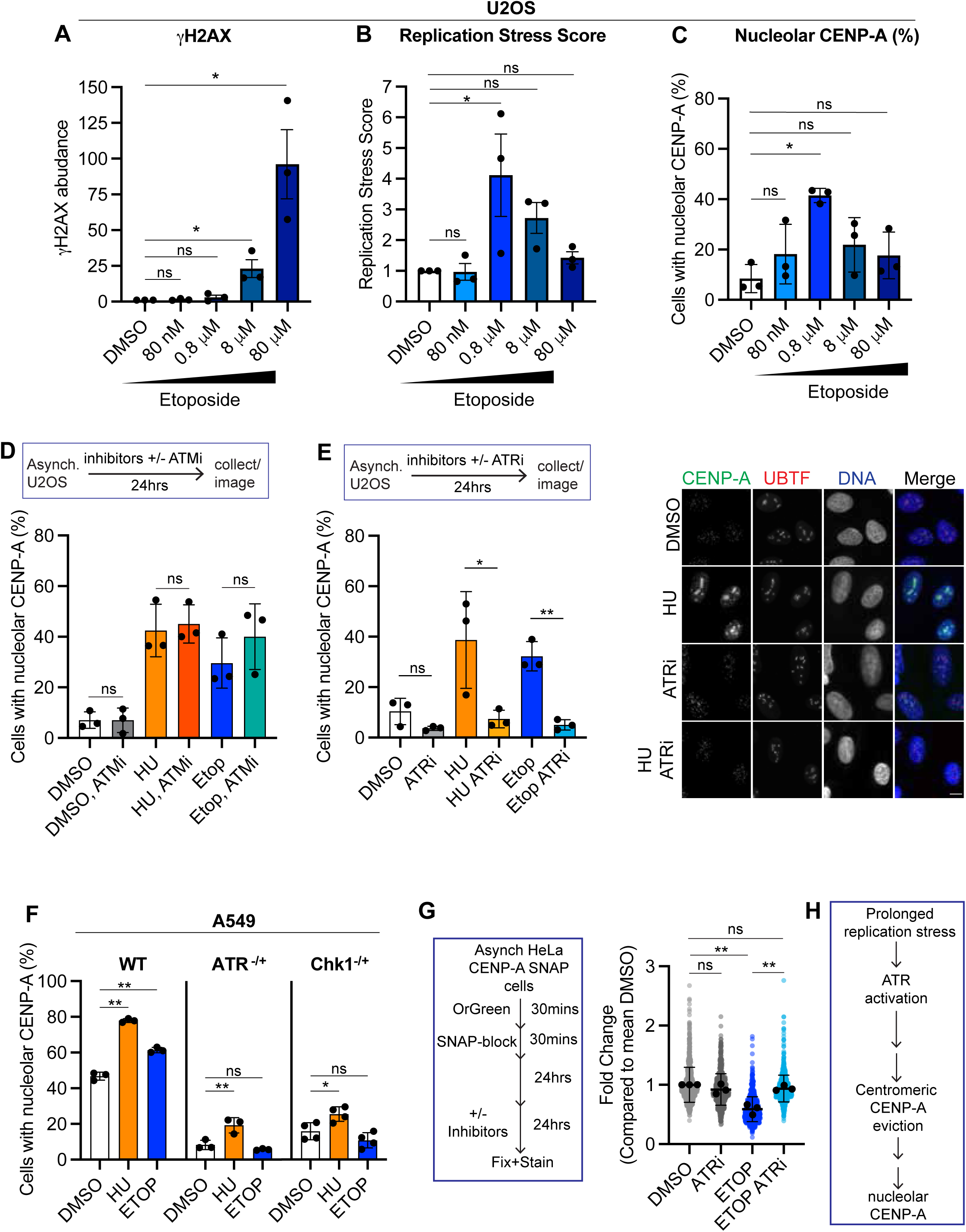
CENP-A relocalization to nucleoli relies on ATR activity. (**A**) Quantification of gH2AX abundance in U2OS cells that were treated with either DMSO or various concentrations of Etoposide for 24 hours. Replicates were normalized to DMSO. (**B**) Quantification of replication stress score of U2OS cells after treatment with Etoposide as in (A). Replication stress score was calculated by multiplying the fold decrease in EdU incorporation in S-phase cells by the percentage of S-phase cells. Data was then normalized to DMSO (**C**). Percentage of cells that contain nucleolar CENP-A when treated as in (A). (**D**) Percentage of cells with nucleolar CENP-A after 24-hour treatment with either DMSO, 10μM KU (ATMi), 1mM hydroxyurea (HU), 3μM Etoposide (ETOP), or in combination. (**E**) Percentage (left) and representative images (right) of cells with nucleolar CENP-A after 24-hour treatment with either DMSO, 10μM AZ20 (ATRi), 1mM hydroxyurea (HU), 3μM Etoposide (ETOP), or in combination. Scale bar =10μm. (**F**) Percentage of A549 parental cells (WT), ATR heterozygous knockout (ATR^-/+^), or Chk1 heterozygous knockout (Chk1^-/+^) cells that were positive for CENP-A localization to nucleoli after DMSO, 1mM hydroxyurea (HU) or 3μM Etoposide (ETOP) treatment. (**G**) Quantification of the intensity of “ancestral” CENP-A SNAP at centromeres in stably expressing HeLa cells after treatment with DMSO, 3μM Etoposide (ETOP), 10μM AZ20 (ATRi) alone or in combination. CENP-A, which was labeled 24 hours prior to drug treatment. Centromeres were marked using ACA. Black points represent the mean replicate averages, and error bars represent the standard deviation. *p<0.5; **p<0.01. Significance was calculated using the Student-T test, where there are two conditions only (D,E), or ANOVA with Dunnett test when comparing multiple conditions to DMSO (A-C, F), or Tukey test when comparing across all conditions (G).

We thus sought to directly test if CENP-A relocalization to nucleoli was dependent on ATR or ATM activity by inhibiting these kinases using small-molecule inhibitors. U2OS cells were treated with ATM inhibitor (10μM, KU-55933) alone or in combination with 1mM HU or 3μM ETOP for 24 hours. We observed that treatment of U2OS cells with the ATM inhibitor did not decrease the propensity of cells having relocalized CENP-A to nucleoli, compared to DMSO-treated cells (**Figure 3D**). Additionally, co-treatment of ATMi did not rescue relocalization of CENP-A to nucleoli in HU or ETOP-treated cells (**Figure 3D**). In contrast, ATR inhibition led to a decrease in CENP-A relocalization when cells were co-treated with either HU or ETOP (**Figure 3E**). We further observed that inhibition of Chk1, ATR’s effector kinase, also led to a decrease in CENP-A relocalization to nucleoli when cells were co-treated with either HU or ETOP (**Figure S4A**).

Consistent with these data, A549 cells that had decreased ATR or Chk1 due to heterozygous knockout (ATR+/-, Chk1+/-) did not exhibit increased rates of CENP-A relocalization to nucleoli after HU treatment (**Figure 3F, Figure S4B**). These data suggest that in response to prolonged DNA damage, human CENP-A relocalization relies on ATR signaling.

Due to the dependence of ATR in human cells, we decided to explore the possibility that CENP-A relocalization to nucleoli might still be ATR-dependent in mouse NIH-3T3 cells during replication stress. NIH-3T3 cells were treated with 1mM HU or 3μM ETOP for 24 hours. There was an increase in the percentage of cells with nucleolar CENP-A in response to ETOP or HU (**Figure S4C, D**). Similarly to previous results, when cells were co-treated with ETOP and ATMi, we observed a small but significant decrease in CENP-A localization to nucleoli (**Figure S4C, D**). However, the decrease was present but not significant in HU-treated cells (**Figure S4C**).

Importantly, treatment with ATMi did not fully rescue the CENP-A phenotype to control levels. In contrast, when NIH-3T3 cells were similarly co-treated with ATRi, we observed a dramatic decrease in the percentage of cells exhibiting nucleolar CENP-A, similar to that observed with DMSO treatment (**Figure S4C, D**). We further observed a similar rescue when NIH-3T3 cells were treated with Chk1i (**Figure S4E**). These data demonstrate that in response to replication stress, in human and mouse cells, ATR signaling is necessary for CENP-A relocalization to the nucleolus.

We hypothesized that this process of CENP-A relocalization to the centromere required two steps. The first is the loss of CENP-A from centromeres, and the second is the relocalization of CENP-A to nucleoli. We sought to determine if ATR prevented CENP-A localization to nucleoli by preventing centromeric loss. To test this, we used CENP-A SNAP-expressing HeLa cells and measured centromeric CENP-A after ETOP treatment (**Figure 3G**). We observed that cells treated with ATRi showed a small decrease in CENP-A SNAP intensity at centromeres (marked by anti-centromere antibody), which was smaller than the decrease observed in cells treated with ETOP (Figure 3G). Importantly, co-treatment of ETOP with ATRi led to a rescue of CENP-A SNAP intensity back to that of ATRi alone-treated cells (**Figure 3G**). These data suggest that ATR promotes CENP-A occupancy loss in response to DNA damage (**Figure 3H**).

### HJURP promotes CENP-A relocalization to nucleoli

Due to the effects observed after ATRi and our previous data^11^, we hypothesized that the H3 histone chaperone DAXX could be involved in CENP-A eviction and/or CENP-A relocalization to nucleoli. We sought to measure the percentage of nucleolar CENP-A-positive A549 cells that were WT for DAXX or were DAXX null using CRISPR knockout (DAXX^-/-^). DAXX^-/-^ A549 cells were WT for ATRX, an additional H3 histone chaperone that cooperates with DAXX, but did have decreased levels of ATRX compared to WT cells (**Figure S5**) ^26^. DMSO-treated DAXX^-/-^ had a lower percentage of cells that were positive for nucleolar CENP-A (∼16%) compared to WT cells (32%). After 24-hour treatment with hydroxyurea (HU), we observed a significant increase in the percentage of cells positive for nucleolar CENP-A, comparable to the increases observed in WT cells (∼283% and 128% increase, respectively) (**Figure 4A, B**). Importantly, these data are vastly different than what we observed when cells were depleted of ATR or inhibited with an ATR inhibitor (**Figure 3E, F**), suggesting that DAXX is not the main regulator of relocalization to the nucleolus in response to sustained replication stress. Consistent with these data, DAXX has been observed to be maintained at PML nuclear bodies and phosphorylated by ATM, where it resides, in response to DNA damage ^27,28^.

**Figure 4:**
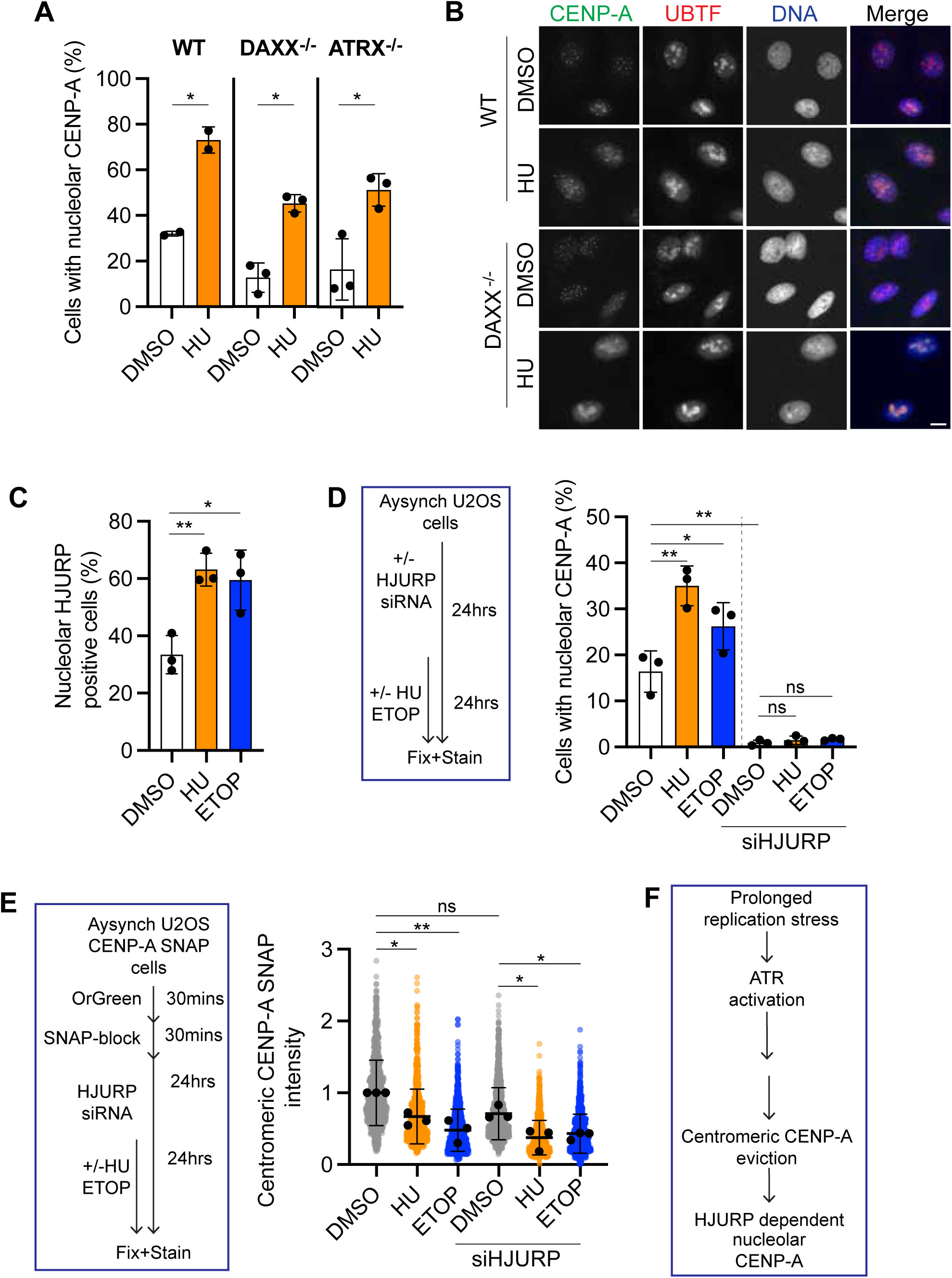
HJURP promotes CENP-A relocalization to nucleoli. (**A**) Percentage of A549 parental (WT), DAXX knockout (DAXX-/-), or ATRX knockout (ATRX-/-) cells that were positive for nucleolar CENP-A after 24-hour treatment with DMSO or HU (1mM hydroxyurea). (**B**) Representative image of WT and DAXX-/- A549 cells that were treated as in (A). Scale bar =10μm. (**C**) Percentage of U2OS cells that exhibited HJURP localization to the nucleolus (marked by UBTF). Positive localization was determined using colocalization analysis and deemed positive if R(ob) was greater than or equal to 0.65. (**D**) Percentage of WT or HJURP-depleted U2OS cells that exhibit CENP-A localization to the nucleolus (marked by UBTF) after 24-hour treatment with DMSO, HU (1mM hydroxyurea), or ETOP (3 μM Etoposide). HJURP depletion was done for a total of 48 hours. (**E**) Quantification of the intensity of “ancestral” CENP-A SNAP at centromeres in stably expressing HeLa cells that were either WT or HJURP-depleted and subsequently treated with DMSO, HU, or ETOP as in (D). Cells were labeled 24 hours prior to HJURP knockdown. Black points represent the mean replicate averages, and error bars represent the standard deviation. *p<0.5; **p<0.01. Significance was calculated using the Student-T test, where there are two conditions only (A), or ANOVA with Dunnett test when comparing multiple conditions to DMSO (B), or Tukey test when comparing across all conditions (D,E).

Similar to DAXX, ATRX has been implicated in CENP-A eviction, especially in response to replication stress in non-cancer cells^17^. Thus, we further assessed the effect of ATRX on CENP-A relocalization in response to sustained replication stress. ATRX -/- cells did not exhibit any significant loss of DAXX compared to WT cells (**Figure S5A**). ATRX -/- cells exhibited a low rate of cells with nucleolar CENP-A in DMSO treatment (∼16%) (**Figure 4A**). 24-hour HU treatment resulted in a significant increase in the percentage of cells positive for nucleolar CENP-A (∼51%), representing a similar percentage increase (∼200%) (**Figure 4A**). Taken together, these data suggest that neither DAXX nor ATRX is involved in CENP-A eviction from the centromere or relocalization to nucleoli in response to DNA damage.

These data led us to interrogate the role of HJURP in this process. HJURP is the canonical CENP-A chaperone and can localize with nascent CENP-A to the nucleolus, especially in S- phase . We observed that there was an increase in HJURP localization to nucleoli after HU or ETOP treatment compared to DMSO treated cells (**Figure 4C**), leading the possibility that indeed HJURP could help translocate CENP-A. To test this directly, we knocked down HJURP using siRNA (HJURPsi) for 24 hours, then treated cells with either DMSO, HU, or ETOP for 24 hours, as in prior experiments, and assessed CENP-A localization to nucleoli (**Figure 4C**). We observed that in HJURPsi-treated cells, there was an almost complete loss of CENP-A localization to nucleoli in DMSO cells compared to HJURP proficient cells (**Figure 4D**).

Moreover, neither HU nor ETOP treatment leads to an increase in CENP-A localization to nucleoli (**Figure 4D**), suggesting that HJURP is necessary for CENP-A relocalization after replication stress. To define if HJURP was necessary for CENP-A eviction from the centromere or CENP-A localization to the nucleolus, we used CENP-A SNAP expressing U2OS cells where we labeled CENP-A SNAP 24 hours prior to HJURP knockdown and subsequent treatment with either DMSO, HU, or ETOP (**Figure 4E**). Consistent with prior observations, we observed that there was a moderate but significant decrease in CENP-A SNAP intensity at centromeres in HJURPsi-treated cells, compared to untreated cells (**Figure 4E**)^29^. However, we observed a further decline in centromeric CENP-A SNAP intensity after treatment with HU or ETOP (**Figure 4E**), demonstrating that loss of HJURP does not prevent CENP-A eviction from the centromere. Taken together, these data suggest that HJURP is necessary for CENP-A relocalization after centromeric eviction during replication stress (**Figure 4F**). However, these data did not answer how ATR in response to replication stress promoted centromeric CENP-A eviction.

### ATR promotes CENP-A eviction via p97

We decided to assess known factors that remove CENP-A from the chromatin. The ATPase, VCP (p97), promotes the removal of CENP-A via the recognition of SUMOylated CCAN components, which facilitates their degradation and destabilizes CENP-A-containing nucleosomes in S-phase ^30^. Thus, we sought to determine if VCP was involved in this CENP-A relocalization pathway that occurs after replication stress by assessing nuclear CENP-A after VCP inhibitor treatment.

We treated U2OS cells with VCP inhibitor (CB-5083) for 24 hours in combination with hydroxyurea (HU) or Etoposide (ETOP) and measured the percentage of cells with nucleolar phenotype (**Figure 5A,B**). Under these conditions, HU and ETOP treatment led to a significant increase in the percentage of cells with nucleolar CENP-A compared to DMSO-treated cells(**Figure 5A**). Co-treatment of p97i in HU-treated cells or ETOP-treated cells decreased the percentage of cells exhibiting nucleolar CENP-A (**Figure 5A**). Importantly, VCPi also led to a decrease in CENP-A localization to nucleoli in NIH-3T3 cells (**Figure S6A**). These data suggest that VCP is involved in the pathway that leads to CENP-A eviction and relocalization to nucleoli including in mouse cells.

**Figure 5:**
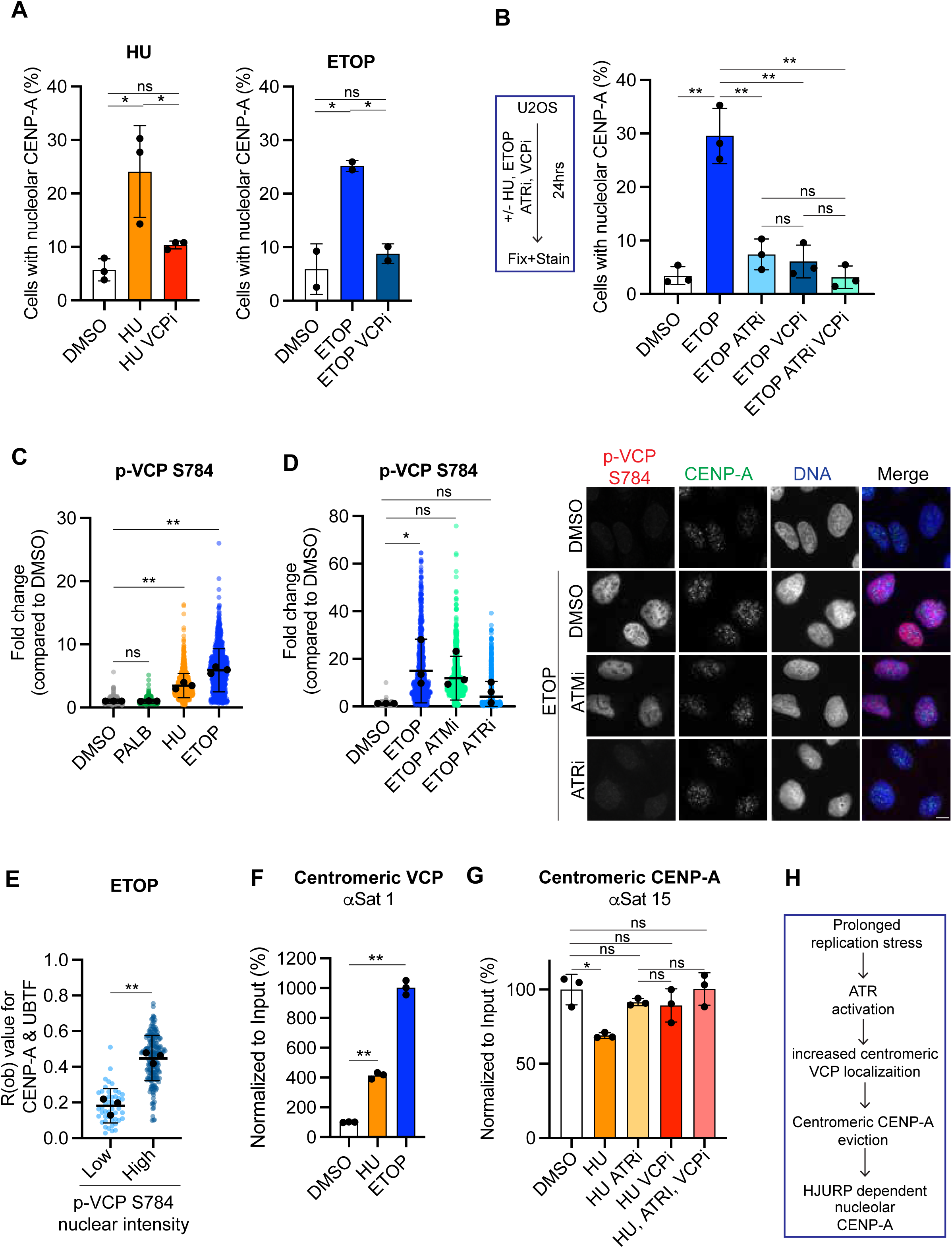
ATR promotes CENP-A eviction via p97. (A) Percent of U2OS cells that exhibit CENP-A localization to the nucleolus (marked by UBTF) treated with DMSO, 1mM hydroxyurea (HU) or co-treated with 1mM hydroxyurea and XμM p97inhibitor-name (HU VCPi) (right) or treated with DMSO, 3μM Etoposide (ETOP) or 3μM Etoposide and XμM p97inhibitor-name (ETOP VCPi). (**B**) Percent of U2OS cells that exhibit CENP-A localization to the nucleolus (marked by UBTF) treated with DMSO 3μM Etoposide (ETOP) or co-treated with 3μM Etoposide (ETOP) and 10 μM AZ20 (ATRi), XμM p97inhibitor-name (VCPi) or both ATRi and VCPi. (**C**) Quantification of nucleolar phosphorylated VCP S784 in U2OS cells after 24hour treatment with DMSO, 1 μM Palbociclib (PALB), 1mM Hydroxyurea (HU), and 3μM Etoposide (ETOP). (**D**) Black points represent the mean replicate averages, and error bars represent the standard deviation. *p<0.5; **p<0.01. Significance was calculated using the Student-T test, where there are two conditions only (E), or ANOVA with Dunnett test when comparing multiple conditions to DMSO (C, D, G), or Tukey test when comparing across all conditions (A,B,F).

Recent evidence demonstrates that VCP is phosphorylated in response to double-stranded breaks at S784, thereby enhancing p97 activity, leading to the possibility that VCP could be regulated by ATR or other DNA damage kinases ^31^.We sought to determine if ATR and VCP were in the same pathway, by treating etoposide treated cells with either ATR inhibitor (ATRi) or VCP inhibitor (VCPi) alone or in combination. We observed that either ATRi or VCPi led to a significant decrease in cells exhibiting nucleolar CENP-A, and that when combined, a slightly lower, but not statistically significant, decrease was observed (**Figure 5B**). These suggest that VCP and ATR are in the same pathway.

We decided to further investigate whether VCP was regulated by ATR by measuring phosphorylation of VCP. We first assessed VCP phosphorylation in the nucleolus using a commercially available antibody against p-VCP S784, the known DNA damage-induced phosphorylation site. Indeed, there was a significant increase in nuclear intensity of p-VCP S784 in cells treated with either HU or ETOP for 24 hours, compared to cells treated with either DMSO or Palbociclib (PALB) (**Figure 5C**). Co-treatment with ATRi in ETOP-treated cells significantly decreased p-VCP S784 to levels comparable to those observed with DMSO treatment (**Figure 5D**). However, inhibition of ATM did not significantly decrease p-VCP S784 nuclear intensity (**Figure 5D**). Due to this striking increase in p-VCP S784 in ETOP treatment (**Figure 5C**), we sought to determine if there was any correlation between p-VCP S784 increase and CENP-A nucleolar localization. Indeed, we observed that cells with high p-VCP S784 nuclear intensity (above 1.5-fold increase compared to the average intensity of DMSO-treated cells) were more likely to have higher nucleolar CENP-A localization (using Rob values) compared to cells with “low p-VCP S784” nuclear intensity in ETOP-treated cells (**Figure 5E**). In DMSO-treated cells, we observed a moderate increase in CENP-A localization to nucleoli, but this was not significant, likely due to the low number of cells with elevated p-VCP S784 nuclear intensity (**Figure S6B**). Consistent with our data in U2OS cells, we also observed a significant increase in p-VCP S784 in mouse NIH3T3 cells in response to HU and ETOP (**Figure S6C**).

Moreover, co-treatment with etoposide (ETOP) and ATRi but not ATMi led to a significant decrease in p-VCP S784 that was comparable to DMSO (**Figure S6D**). Taken together, these data suggest that in response to prolonged replication stress, VCP is regulated in an ATR- dependent manner.

We then sought to specifically define the role of VCP and how ATR regulates its function in this nuclear CENP-A relocalization pathway. Due to its known function in promoting CENP-A eviction from chromatin, we hypothesized that VCP was crucial in removing CENP-A from the centromere. We also observed that the occupancy of CENP-C and CENP-T, components of the CCAN and known substrates of VCP in S-phase,^30^ were significantly decreased in response to HU (**Figure S6D, E-F**), supporting our observation that CENP-A loss was VCP(p97) dependent. Consistent with these data, we observed that VCP was highly enriched at centromeric chromatin in response to HU and ETOP (**Figure 5F, Figure S6G**). These data led us to test the role of VCP and ATR on CENP-A occupancy directly. Using ChIP, we observed that inhibition of VCP or ATRi alone after HU treatment led to a rescue of CENP-A occupancy at centromeres (**Figure 5G, Figure S6H**). Moreover, in ATRi combination with VCPi led to a similar rescue of CENP-A occupancy at centromeres (**Figure 5G, Figure S6H**). Taken together, these data suggest that ATR promotes VCP localization to centromeres in response to prolonged replication stress, which leads to CENP-A eviction from the centromere (**Figure 5H**).

### ATR-dependent loss of CENP-A is responsible for increased acentric fragments after replication stress

Increased acentric fragment missegregation has been widely observed as a consequence of replication stress^5^. We hypothesized that the cause of these acentric fragments could in part be due to the ATR-dependent loss of CENP-A from centromeric chromatin. We first sought to determine if CENP-A remained low once cells began cycling again. We treated CENP-A SNAP-expressing HeLa cells with either Palbociclib (PALB) or hydroxyurea (HU) for 24 hours and subsequently released them into fresh media for an additional 24 hours (**Figure 6A**). We analyzed mitotic cells in these populations and measured SNAP-CENP-A that was labeled 24 hours prior to any treatment, as before. Mitotic cells that were previously treated with HU exhibited lower CENP-A SNAP intensity at centromeres compared to palbociclib-released cells (**Figure 6A**), suggesting that the loss of ancestral CENP-A from centromeres is not redeposited after the resolution of replication stress and cell cycle restart.

**Figure 6:**
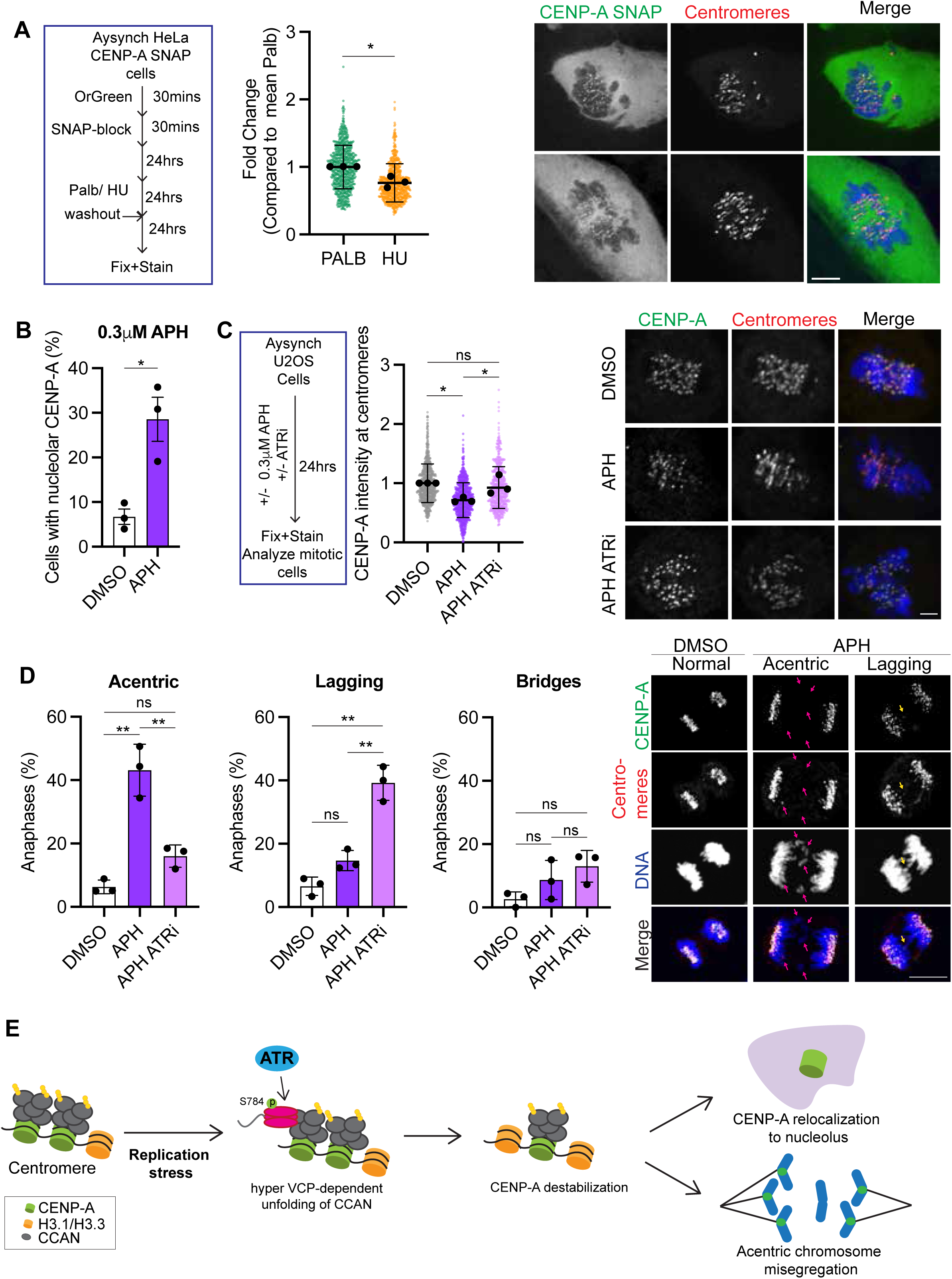
ATR-dependent loss of CENP-A is responsible for increased acentric fragments after replication stress. Schema and quantification (left) and representative images (right) of the intensity of “ancestral” CENP-A SNAP at centromeres in stably expressing HeLa cells after treatment with 0.5 μM Palbociclib or 1mM Hydroxyurea (HU) for 24 hours and subsequent washout and cell cycle entry exhibit. Centromeres were marked using ACA. Scale bar = 5μm. (**B**) Percent of U2OS cells that exhibit CENP-A localization to the nucleolus (marked by UBTF) treated with DMSO or 3 μM Aphidicolin (APH) for 24 hours. (**C**) Schema and quantification (left) and representative images (right) of CENP-A intensity at centromeres in mitotic cells. Cells were treated with either DMSO, 3μM Aphidicolin (APH), or 3μM Aphidicolin and 1μM AZ20 (APH ATRi). Scale bar =5μm. (**D**) Percentage of anaphases that exhibit acentric (chromosome missegregation that is absent of CENP-A), lagging (a chromosome missegregation where CENP-A is present), or chromosome bridges (DNA connecting both masses) in U2OS cells treated as in (D). Representative image (right) of anaphases stained with CENP-A (green), anti-centromere antibody (red) and DAPI with no chromosome segregation defects or acentric or lagging chromosomes arising in APH-treated cells quantified on the left. Acentric fragments are noted with a pink arrow, while lagging chromosomes are noted with a yellow arrow. Images are maximal projections of Z-stacks. Scale bar = 10μm. (**E**) Model demonstrating that prolonged replication stress leads to CENP-A dependent phosphorylation of VCP at S784, promoting VCP localization to centromeres, which destabilizes CENP-A containing nucleosomes. Evicted CENP-A is subsequently relocalizes to nucleoli in an HJURP dependent manner. Loss of centromeric CENP-A persists through mitosis and leads to the increase in acentric chromosome segregation observed. Black points represent the mean replicate averages, and error bars represent standard deviation. *p<0.5; **p<0.01. Significance was calculated using the Student-T test, where there are two conditions only (A,B), or ANOVA with Tukey test when comparing across all conditions (C,D).

We then sought to test our hypothesis by inducing low-grade replication stress that did not lead to a complete cell cycle arrest. We decided to use 0.3mM aphidicolin (APH) treatment, which we and others have demonstrated allows cells to continue through mitosis with high missegregation rates ^4,32^. Under these conditions, we observed a significant increase (∼200% increase) in the percentage of cells exhibiting nucleolar CENP-A (**Figure 6B**). We then sought to determine if mitotic cells treated with APH had decreased CENP-A occupancy at centromeres by measuring CENP-A intensity. Indeed, APH led to a decrease in centromeric CENP-A in mitotic cells (**Figure 6C**). This was rescued by co-treatment with a low dose of ATRi (1uM AZ20) (**Figure 6C**). We then measured missegregation errors in mitosis under these conditions. We observed that acentric missegregation events, defined as chromatids outside chromatid planes in anaphase that lacked CENP-A, were highly elevated in APH-treated cells, but were decreased significantly in cells that were co-treated with APH and ATRi (**Figure 6D**). Conversely, APH did not lead to significantly increased lagging chromosome missegregation events, defined as chromatids that were outside the chromosome planes in anaphase that contained CENP-A (**Figure 6D**). Co-treatment of APH and ATRi resulted in a dramatic and significant increase in lagging chromosomes (**Figure 6D**). The increase in lagging chromosomes is likely due, in part, to the mitotic function of ATR ^10^. Importantly, we observed a mild and similar increase in chromosome bridges, where chromosomes fuse together, leading to missegregation, in both APH alone and in APH and ATRi co-treated cells (**Figure 6D**). Taken together, these data suggest that replication stress leads to an ATR-dependent loss of CENP-A, which in turn leads to acentric segregation defects in mitosis (**Figure 6E**).

## DISCUSSION

Our study uncovers a previously unidentified role for ATR in regulating centromere identity in response to replication stress. We show that ATR promotes eviction of the centromeric histone H3 variant CENP-A, followed by CENP-A nucleolar relocalization – an HJURP dependent process. CENP-A eviction occurs in an ATR and AAA+ ATPase VCP (p97) dependent manner. Partial loss of CENP-A persists after cell-cycle re-entry post replication stress-dependent S-phase arrest, and lasts into mitosis. Thus, replication stress correlates with increased acentric chromosome missegregation in mitosis. These findings identify ATR as a direct regulator of centromere composition and establish a mechanistic link between replication stress and genome instability.

ATR is the master kinase of the DNA damage response pathway, and its functions in stabilizing stalled forks and activating the replication checkpoint have been well studied. Our previous study demonstrated that inhibiting ATR is also deleterious for centromere identity via DAXX-dependent CENP-A eviction^11^. Our new results demonstrate that high ATR activity due to DNA damage is also deleterious to centromeres, again leading to CENP-A eviction. These results demonstrate that centromeres are exquisitely sensitive to ATR activity. Moreover, the activity from ATR does not have to be local. While its most established role works at the local level at single-stranded DNA lesions, our data suggest that CENP-A eviction occurs as a consequence of a more global effect of ATR activity. We did not identify direct ATR activation at centromeres via increased single-stranded DNA, which we probed using Rad51, which binds directly to single-stranded DNA and is an early step in DNA repair (**Fig. S1F**). Importantly, unlike γH2AX, which is not always observed even if DNA damage is present, Rad51 has been observed at centromeres^6,33^.

Our results further suggest that ATR-dependent CENP-A eviction may contribute to the genome instability observed in many cancer cells, that exhibit replication stress. ATR inhibitors are currently in clinical trials. These inhibitors may therefore alter centromere identity in addition to checkpoint and repair pathways. Depending on context, they may stabilize centromeres (as we observed in cancer cells with replication stress), but in non-cancer cells or cancer cells that do not are not undergoing long-term replication stress, ATR inhibitor may promote centromere dysfunction, as noted in our previous work^11^. These are important implications for therapeutic strategies that involve using ATR inhibitor.

We identify that ATR promotes the activity of VCP (p97), which destabilizes CENP-A nucleosomes. VCP has been implicated in chromatin remodeling and extraction of misfolded proteins after DNA damage^34^. In S-phase, VCP can destabilize CENP-A by promoting the unfolding of Constitutive Centromere Associated Network (CCAN) components ^30^. Our data suggest that in response to prolonged replication stress, ATR phosphorylates VCP, leading to increased nuclear localization and binding to centromeres (**Figure 5D, 5F**). If ATR or other downstream components also promote VCP activity and its localization, or just this increased localization to centromeres is sufficient to increase CENP-A stabilization and eviction, is unclear and remains a question that should be answered in the future. Our data in mouse cells, also shows that the ATR-VCP pathway is the predominant pathway by which CENP-A eviction occurs in response to DNA damage. While ATM may be an important player in CENP-A eviction in response to etoposide in mouse cells via direct phosphorylation, its inhibition after prolonged treatment with hydroxyurea or etoposide did not relieve CENP-A eviction and translocation to the nucleolus. This is in stark contrast with previous work using NIH3T3 cells. However, the concentration of etoposide used in those studies may have led to a different arrest. This leaves the possibility that cells may rely on ATR and ATM at different cell cycle stages to get the same result of CENP-A eviction.

Following eviction, CENP-A relocalizes to nucleoli in a process dependent on its histone chaperone, HJURP, but not the H3.3 histone chaperones, DAXX or ATRX (**Figure 4**). During normal cell cycle progression, HJURP is found in the nucleolus with pre-nucleosomal CENP-A^14,15^. It is possible that evicted CENP-A is chaperoned to the nucleolus by HJURP to be stored in nucleoli in a similar manner. This may be a necessary step to prevent ectopic CENP-A deposition at DNA-damaged sites^35^. It is unclear if additional partners or modifications on CENP-A are necessary for CENP-A localization to the nucleolus to occur. Moreover, it is also possible CENP-A localization to the nucleolus is important for nucleolar function. We do not anticipate that CENP-A is incorporating into the DNA, because to our fractionation results show that CENP-A is removed from the chromatin bound pool (**Figure 1C**). However, CENP-A is known to interact with NPM1 and possibly interacts with other nucleolar proteins, leaving the possibility that CENP-A localization is indeed necessary for proper nucleolar function^36^. These findings suggest that HJURP functions not only as a centromeric chaperone but also as a stress-responsive factor that redistributes CENP-A to nucleoli.

Recent work from the Foltz lab demonstrates that replication stress in RPE-1 cells, non-cancer cells, leads to an ATRX-dependent loss of CENP-A from centromeres after aphidicolin ^17^. In lung adenocarcinoma, A549 cells, which lack either DAXX or ATRX, did not rescue nucleolar CENP-A relocalization in response to hydroxyurea (**Figure 4A**). Taken together, this suggests that cancer cells and normal cells may rely on DAXX and ATRX differently, possibly due to increased CENP-A expression in cancer cells. This may represent an important contrast between healthy and cancer cells, which could be further exploited.

As expected, ATR-dependent CENP-A eviction from the centromere is not reversible in G2, likely due to the normal regulation of CENP-A deposition which occurs only in early G1. This leaves cells to have to contend with decreased CENP-A at centromeres in mitosis. Likely, centromeric replacement of CENP-A occurs, as we continue to observe similar levels of H2A at centromeres in HU-treated cells compared to palbociclib-treated cells (**Figure S2F**). Similarly to what we previously observed, the loss of CENP-A and possible replacement with H3.3 or other histones is likely the cause of the increased rate of acentric chromosomes observed after mild replication stress (**Figure 6D**)^11^. These results provide a direct mechanistic connection between replication stress and chromosome segregation errors, thereby linking centromere dysfunction to chromosomal instability. The persistence of eviction suggests that transient replication stress has a long-term impact on centromere identity, with lasting consequences for genome stability. Why CENP-A eviction seems to be so sensitive to ATR activity remains unclear. It is possible that CENP-A eviction from centromeres may act as a backup mechanism to cause cell death or cell cycle arrest post-high replication stress. Unlike double-stranded breaks, cells can enter mitosis and continue cycling with a moderate abundance of single-stranded lesions^37,38^. Thus, it is possible that cells employ additional layers of protection to ensure that cell cycle arrest or removal from the population occurs via autonomous or non-autonomous mechanisms after replication stress. Chromosome missegregation events are known to lead to cell cycle arrest in the G1 and S phases, halting the cell cycle. While missegregation of chromosomes can lead to the formation of micronuclei, which can, in turn, signal the innate immune system to remove these cells. Thus, ATR-dependent CENP-A loss and centromere disassembly may be a mechanism to ensure cell cycle arrest or death if the cell is able to overcome the replication stress.

In conclusion, we propose a model in which ATR links replication stress to genome instability by evicting CENP-A from centromeres and directing it to nucleoli. This pathway connects replication stress signaling directly to centromere identity, providing a mechanism by which stress promotes segregation errors and chromosomal instability.

## ACKNOWLEDGEMENTS

This work is supported by R35 GM150648 (LK), R35 GM150645 (KS). We thank Dr. Lars Jansen for generously providing the CENP-A-SNAP expressing HeLa cells. We thank the members of the Kabeche Lab and Yale’s Cancer Biology Institute for their critical feedback on the manuscript.

## AUTHOR CONTRIBUTIONS

I.T., D.O., H.L., K.S., and L.K. designed the study. I.T., D.O., H.L., R.S., E.Z., C.B., K.S. and L.K. performed the experiments and analyses. H.L., D.O., and L.K. prepared the manuscript with contributions from all the authors.

## COMPETING INTERESTS

The authors declare no competing interests.

## SUPPLEMENTAL FIGURE LEGENDS

**Figure S1 (related to Figure 1):** (**A**) Western blot of U2OS cells treated with DMSO, 1 mM hydroxyurea (HU), or 3 μM etoposide (ETOP) for 24 hr and probed for γH2AX and p53. (**B**) EdU incorporation plotted against DAPI intensity of U2OS cells treated for 24 hr with DMSO, 1 mM HU, or 3 μM ETOP. (**C**) Western blot of U2OS cells treated with DMSO or 0.5 μM palbociclib (PALB) for 24 hr and probed for γH2AX and p53. (**D**) Western blot of A549 cells treated with DMSO, 1 mM HU, or 3 μM ETOP for 24 hr and probed for γH2AX and p53. (**E**) Western blot of HeLa cells treated with DMSO, 1 mM HU, or 3 μM ETOP for 24 hr and probed for γH2AX. (**F**) Quantitative PCR of histone H2A chromatin immunoprecipitation from U2OS cells treated for 24 hr with DMSO, 1 mM HU, or 0.5 μM PALB, using primers corresponding to centromeric α- satellite regions of chromosomes 1 (αSat1), 4 (αSat4), and 15 (αSat15). Black points represent independent biological replicates, bars represent the mean, and error bars indicate SD. (**G**) Quantification of Rad51 intensity at centromeres in U2OS cells treated with DMSO, 1 mM HU, or 3 μM ETOP for 24 hr, measured by immunofluorescence. Black points represent independent biological replicates, bars represent the mean, and error bars indicate SD. Statistical significance was calculated using one-way ANOVA with Dunnett’s test when comparing multiple conditions to DMSO (F), or Tukey’s test when comparing across all conditions (E). *p < 0.05; **p < 0.01; ns, not significant.

**Figure S2 (related to Figure 2):** (**A**–**B**) Representative images of U2OS cells treated with 3 μM etoposide (ETOP) or 0.5 μM palbociclib (PALB) for 24 hr, stained for CENP-A (green), UBTF (red), and DNA (blue). Line scans show fluorescence intensity across the indicated axis, and R(ob) values are reported for each condition. (**C**–**D**) Quantification of R(ob) values measuring colocalization in U2OS cells treated with DMSO, 1 mM hydroxyurea (HU), or 3 μM ETOP (C), or with DMSO or 0.5 μM PALB (D). Each point represents a single nucleus. (**E**) Percentage of U2OS cells with nucleolar CENP-A after 24 hr treatment with DMSO or HU, using a different mouse monoclonal antibody against CENP-A. Representative images are shown on the right. (**F**) Percentage of U2OS cells with nucleolar CENP-A after 24 hr treatment with DMSO, HU, or ETOP in the presence of 1 μM BMH-21 to disrupt nucleoli. Representative images (right) show disintegration of UBTF foci upon BMH-21 treatment. (**G**) Percentage of U2OS cells with nucleolar CENP-A after exposure to 1.4 Gy ionizing radiation (IR) and fixation 24 hr later. (**H**) Quantification of cells positive for nascent CENP-A, labeled by pulsing SNAP after treatment with DMSO, 1 mM HU, or 3 μM ETOP for 24 hr and prior to fixation. Black points represent independent biological replicates, bars represent the mean, and error bars indicate SD. Statistical significance was calculated using Student’s t-test for pairwise comparisons (D, E, G) or Tukey’s test when comparing across all conditions (C, F). *p < 0.05; **p < 0.01; ns, not significant.

**Figure S3 (related to Figure 3):** (**A**) Western blots of U2OS cells treated for 24 hr with DMSO or decreasing concentrations of etoposide (80 μM, 8 μM, 0.8 μM, 80 nM). Blots were probed for γH2AX and CENP-A (Rb). (**B**) Quantification of EdU incorporation in S-phase U2OS cells normalized to DMSO after etoposide treatment of increasing concentration (80 nM, 0.8 μM, 8 μM, 80 μM). Black points represent independent biological replicates; error bars show SD. (**C**) Representative images of U2OS cells treated with DMSO or the indicated concentrations of etoposide for 24 hr and stained for CENP-A (green), UBTF (red), and DNA (blue). Merged images are shown on the right. Scale bar =10μm. Statistical significance was calculated using one-way ANOVA with Dunnett’s test when comparing multiple conditions to DMSO (B). *p < 0.05; ns, not significant.

**Figure S4 (related to Figure 3):** (**A**) Quantification of U2OS cells with nucleolar CENP-A after 24 hr treatment with DMSO, 1 mM hydroxyurea (HU), or 3 µM etoposide (ETOP), in the presence or absence of 1 μM Chk1 inhibitor (Chk1i). Black points represent independent biological replicates, bars show mean ± SD. (**B**) Western blots of WT, Chk1+/–, and ATR+/– heterozygous knockout A549 cells probed for ATR and Chk1. (**C**) Quantification of NIH-3T3 cells with nucleolar CENP-A after 24 hr treatment with DMSO, 1 mM HU, or 3 µM ETOP in the presence or absence of 10 μM ATM inhibitor (ATMi) or 1 μM ATR inhibitor (ATRi). Black points represent independent biological replicates, bars show mean ± SD. (**D**) Representative images of NIH-3T3 cells stained for CENP-A (green), UBTF (red), and DNA (blue) as treated in (C). Scale bar =5μm. (**E**) Quantification of NIH-3T3 cells with nucleolar CENP-A after 24 hr treatment with DMSO or 3 µM ETOP, with or without 1 µM Chk1i. Black points represent independent biological replicates, bars show mean ± SD. Statistical significance was calculated using Student’s t-test for pairwise comparisons (A), one-way ANOVA with Dunnett’s test when comparing multiple conditions to DMSO (A, C), or Tukey’s test when comparing across all conditions (C). *p < 0.05; **p < 0.01; ns, not significant.

**Figure S5 (related to Figure 4):** Western blots of WT, ATRX-/–, and DAXX-/– homozygous knockout A549 cells probed for DAXX and ATRX.

**Figure S6 (related to Figure 5):** (**A**) Percentage of NIH-3T3 cells with nucleolar CENP-A after 24 hr treatment with DMSO, 3 µM etoposide (ETOP), or ETOP in combination with 1 µM VCP inhibitor (VCPi). Black points represent independent biological replicates, bars show mean ± SD. (**B**) Quantification of R(ob) values for CENP-A and UBTF colocalization in U2OS cells treated with DMSO, grouped by low or high nuclear intensity of p-VCP S784. (**C**) Quantification of p-VCP S784 nuclear intensity in NIH-3T3 cells treated with DMSO, 1 mM hydroxyurea (HU), or 3 µM ETOP for 24 hr, normalized to DMSO. Each point represents a single nucleus. (**D**) Quantification (left) and representative images (right) of p-VCP S784 nuclear intensity in NIH- 3T3 cells treated with DMSO or 3 µM ETOP for 24 hr in the presence or absence of 10 µM ATM inhibitor (ATMi) or 10 µM ATR inhibitor (ATRi), normalized to DMSO. Each point represents a single nucleus. Cells were stained for CENP-A (green), p-VCP S784 (red), and DNA (blue). Scale bar = 10µm. (**E-F**) Quantitative PCR of centromeric chromatin immunoprecipitation from U2OS cells treated with DMSO, 1 mM HU, or 3 µM ETOP for 24 hr using antibodies against CENP-C (E) or CENP-T (F). Primers targeted centromeric α-satellite DNA of chromosomes 4 (αSat4) and 15 (αSat15). Black points represent independent biological replicates, bars represent the mean, and error bars indicate SD. (**G**) Quantitative PCR of centromeric chromatin immunoprecipitation from U2OS cells treated with DMSO, 1 mM HU, or 3 µM ETOP for 24 hrs using antibodies against VCP. (**H**) Statistical significance was calculated using Student’s t-test for pairwise comparisons (B,G), one-way ANOVA with Dunnett’s test when comparing multiple conditions to DMSO (C-F,G) and with Tukey’s test when comparing across all conditions (A,H). *p < 0.05; **p < 0.01; ns, not significant.

## MATERIALS AND METHODS

### Cell Culture

RPE-1, U2OS, and HeLa cells were grown in Dulbecco’s Modified Eagle’s medium (Corning, 10-013-CV) supplied with 10% Fetal Bovine Serum (FBS; Corning, 35-011-CV) and 1% Penicillin Streptomycin (P/S; Gibco, 15140-122). Cal27 and Detroit562 were grown in RPMI (Corning, 10-041-CV). A549 wild-type, A549 ATR^+/-^ A549 Chk1^+/-^, A549 DAXX^-/-^, A549 ATRX^-/-^ were grown in Ham’s F-12K Kaighn’s Modification (Gibco, 21127-022) medium supplied with 10% FBS and 1% P/S. All cells were cultured at 37^°^C with 5% CO_2_. For passage, cells were washed 1X with Dulbecco’s Phosphate-Buffered Saline (DPBS; Gibco, 14190-144) and incubated with 0.05% Trypsin-EDTA (Gibco, 25300-054) for 10 minutes at 37^°^C for adherent cells to detach.

### Inhibitors and drug treatments

Small molecule inhibitors used for this study were: ATRi (10μM, AZ-20, Selleckchem), ATM (10 μM KU-55933), Chk1i (2μM MK 8776, Selleckchem), Etoposide (various concentrations, Selleckchem), Palbociclib (1μM, Selleckchem), Aphidicolin (1μM, Selleckchem), Cisplatin (1 μM, Selleckchem), VCPi (1μM CB-5083, Selleckchem).

For all experiments, inhibitors were treated by adding a 1000X stock of inhibitor directly to the plate or wells containing cells at 1:1000 to achieve the final concentration indicated above.

### Antibodies used

Antibodies used:
| REAGENT or RESOURCE | SOURCE | IDENTIFIER |
| --- | --- | --- |
| **Antibodies** |  |  |
| Rabbit Polyclonal Anti-CENP-A | Cell Signaling Technology | 2186 |
| Mouse Monoclonal Anti-CENP-A | Invitrogen | MA1-20832 |
| Rabbit Monoclonal Anti-CENP-A | Invitrogen | MA5-14829 |
| Human anti-centromere antisera (ACA) | Antibodies Inc. | 15-234 |
| Rabbit Polyclonal Anti-DAXX | Invitrogen | PA5-82362 |
| Rabbit Polyclonal Anti-ATRX | Invitrogen | PA5-51656 |
| Rabbit Polyclonal Anti-UBTF | Abnova | H00007343-M01 |
| Rabbit Polyclonal Anti-CENP-C | Invitrogen | MA5-34742 - |
| Rabbit Polyclonal Anti-CENP-T | Invitrogen | PA5-83540 |
| Rabbit Monoclonal Anti-p-VCP S784 | Millipore Sigma | MABS2250 |
| Rabbit Monoclonal Anti-VCP | Abcam | ab109240 |
| Rabbit Monoclonal Anti-Rad51 | Abcam | AB133534 |
| Mouse Monoclonal Anti-Chk1 | Cell Signaling Technology | 2360 |
| Rabbit phospho-p53 (Ser20) | Cell Signaling Technology | 9287 |
| Mouse Monoclonal Anti-Lamin A/C | Cell Signaling Technology | 4777 |
| Mouse Monoclonal Anti-ATR | Novus Biologicals | NB100-308 |
| Mouse Monoclonal Anti-GAPDH-HRP | Santa Cruz Biotechnology | Sc-47724 HRP |
| Normal Rabbit IgG | Cell Signaling Technology | 2729 |
| Rabbit Polyclonal Anti-Histone H2A-ChIP Grade | Abcam | ab18255 |
| Mouse Monoclonal Anti-gamma H2AX [pSer139] | Novus Biologicals | NB100-74435 |
| AffiniPure Goat Anti-Rabbit IgG H+L (Alexa Fluor® 488) | Jackson ImmunoResearch | 111-545 |
| AffiniPure Goat Anti-Rabbit IgG H+L (Alexa Fluor® 594) | Jackson ImmunoResearch | 111-585 |
| AffiniPure Goat Anti-Mouse IgG H+L (Alexa Fluor® 488) | Jackson ImmunoResearch | 115-545 |
| AffiniPure Goat Anti-Mouse IgG H+L (Alexa Fluor® 594) | Jackson ImmunoResearch | 115-585 |
| AffiniPure Goat Anti-Mouse IgG H+L (Alexa Fluor® 488) | Jackson ImmunoResearch | 115-545 |
| AffiniPure Goat Anti-Mouse IgG H+L (Alexa Fluor® 680) | Jackson ImmunoResearch | 115-625 |
| AffiniPure Donkey Anti-Human IgG H+L (Alexa Fluor® 680) | Jackson ImmunoResearch | 709-625 |
| AffiniPure Donkey Anti-Human IgG H+L (Alexa Fluor® 594) | Jackson ImmunoResearch | 709-585 |
| Goat anti-Mouse IgG (H+L) Secondary Antibody, HRP | Invitrogen | 62-6520 |
| Goat anti-Rabbit IgG (H+L) Secondary Antibody, HRP | Invitrogen | 31460 |

#### Immunoblotting

Cells were cultured in 6-well tissue culture plates until confluency was reached. For collection, they were detached using 500 µL of 0.05% Trypsin-EDTA (Gibco) at 37 °C for 10 minutes, then transferred to Eppendorf tubes. The trypsinized cells were centrifuged at 400 × g for 3 minutes, washed with PBS, and resuspended in 50 µL PBS and 50 µL of 2X SDS Sample Buffer (0.1 M Tris-HCl, 4% SDS, 3 mM bromophenol blue, 2 M glycerol, 5% β-mercaptoethanol). Samples were incubated at 95 °C for 15 minutes and lysed using insulin syringes. Proteins were separated on a 7.5% SDS-PAGE gel at 100 V until optimal resolution was achieved and then transferred to 0.2 mM nitrocellulose membranes using the Trans-Blot Turbo Transfer System (Bio-Rad). Membranes were stained with Ponceau S solution (0.5% w/v Ponceau S, 1% glacial acetic acid) for 5 minutes to confirm equal loading, then washed with 1X TBS-T (Tris-buffered saline with 0.1% Tween 20). Membranes were blocked in TBS-T containing 5% BSA for 1 hour at room temperature. Primary antibodies were diluted 1:1000 in blocking solution and incubated with the membranes overnight at 4 °C. The membranes were washed three times for 5 minutes each with TBS-T, followed by incubation with HRP-conjugated secondary antibodies (Invitrogen) diluted 1:1000 in blocking solution for 1 hour at room temperature. After three additional washes, the membranes were incubated with Clarity Western ECL substrate (Bio-Rad) for 5 minutes and imaged using the ChemiDoc MP Imaging System (Bio-Rad).

### Immunofluorescence microscopy

Cells incubated on 18 mm coverslips in 12-well plates were treated with the indicated drugs or reagents, followed by fixation with 3.5% paraformaldehyde for 15 minutes, and then permeabilization using phosphate-buffered saline (PBS) containing 1% Triton X-100 for 10 minutes. Blocking was carried out in PBS with 1% bovine serum albumin (BSA) for 30 minutes. Coverslips were incubated overnight at 4 °C with primary antibodies diluted in the blocking solution. The following day, cells were washed with PBS three times for 5 minutes, and secondary antibodies conjugated to fluorophores, diluted in blocking solution, were then applied along with 1 µg/mL DAPI stain, and incubation proceeded for 1 hour at room temperature (25 °C). After three additional washes in PBS for 5 minutes, coverslips were mounted onto glass slides using ProLong Gold Antifade (Invitrogen) and allowed to dry overnight. Image acquisition was performed using a Nikon ECLIPSE Ti2 widefield epi-fluorescence microscope or on a Nikon ECLIPSE Ti2 W2 spinning disk confocal microscope equipped with a Hamamatsu Fusion sCMOS camera. Images were acquired with NIS-Elements (Nikon), analyzed in FIJI (ImageJ), and processed using Adobe Photoshop.

### EdU treatment and labeling

Newly synthesized DNA was labeled using the Click-&-Go EdU Cell Proliferation Assay Kit (Click Chemistry Tools) and Azide 647. Briefly, asynchronous cells grown on coverslips were treated with 10 mM 5-ethynyl-2′-deoxyuridine (EdU) for 30 minutes. Following treatment, coverslips were processed for immunofluorescence microscopy as described above, with one additional step: after permeabilization with PBS containing 1% Triton X-100, the Click reaction was performed according to the manufacturer’s instructions prior to blocking and incubation with primary antibodies.

Replication stress score were calculated by multiplying the fold-decrease in S-phase EdU incorporation by the percentage of cells in S-phase. (>400 S-phase cells per condition across 3 biological replicates).

### Scoring mitotic defects

All mitotic defects were scored on fixed and stained coverslips. Anaphases were detected using DAPI, whereby there were two chromosome masses and chromosomes were still condensed. Acentric segregation defects were called when a whole chromosome or chromosome fragment was in the middle of two chromosome masses but did not contain CENP-A and was visibly separated from either chromosome mass. These fragments may or may not have been positive for ACA (anticentromere antibody). Lagging chromosomes were called when there was a DNA fragment (DAPI) that was also positive for CENP-A and ACA and was again located between chromosome masses. The DAPI signal that was continuous between the segregating masses, regardless of the ACA signal, was called a chromosome bridge.

### Nuclear intensity quantification

To quantify nuclear fluorescence intensity, single-stack images were acquired at 60X magnification and analyzed in FIJI (ImageJ) using a custom macro. Briefly, nuclei were segmented from the DAPI channel by applying the Huang automatic thresholding algorithm, followed by shape smoothing. Interphase nuclei were identified using the Analyze Particles function with size and circularity thresholds. Nuclei lacking valid ROIs were excluded from further analysis. For each identified ROI, background-subtracted measurements were obtained from both the DAPI channel and the protein-specific channel of interest (e.g., pVCP). For each nucleus, the area, mean intensity, and integrated density were recorded. Results were exported as CSV files for each image and subsequently processed in R. All images across treatment conditions and biological replicates were batch-processed with identical thresholds and measurement parameters.

### Colocalization

To quantify colocalization of CENP-A and UBTF, single-stack images were acquired at 60X magnification and analyzed in FIJI (ImageJ) using a custom macro. Briefly, DAPI masks were generated by thresholding the DAPI channel using the Huang algorithm. Interphase nuclei were segmented and added to the ROI Manager using the Analyze Particles function with specified size and circularity thresholds. An optional threshold based on the maximum intensity of the CENP-A channel was applied at this step to exclude cells lacking specific features (e.g., SNAP, FLAG, HA). For the remaining nuclei, the Colocalization Threshold plugin was used to calculate

Pearson’s coefficients between the CENP-A and UBTF channels within each ROI, following background subtraction. For each image, results were saved as a single CSV file for downstream processing in R. All measurements were batch-processed across different treatment conditions and biological replicates, with all thresholds and analysis parameters held constant within each experiment.

### Cell Cycle Sorting

DAPI and EdU intensity of nuclei were used to classify unsynchronized cell populations into different cell cycle phases. Single-stack images were captured at 20X magnification and processed in FIJI (ImageJ) using a custom macro. The same workflow described for colocalization analysis was applied up to the Colocalization Threshold step, at which point DAPI and EdU intensity for each nucleus were recorded instead. Specifically, nuclei were segmented by thresholding the DAPI channel using the Huang algorithm, followed by binary conversion and particle analysis. ROIs corresponding to nuclei were then used to quantify background-subtracted intensity values for both DAPI and EdU-647 channels. EdU intensity was log-transformed and plotted against DAPI integrated intensity in GraphPad Prism. Based on the clustering of the population, nuclei were grouped into G1, S, or G2 phases.

### Centromeric SNAP quantification

Asynchronous U2OS or HeLa cells stably expressing CENP-A-SNAP were plated and treated with SNAP-Cell substrates TMR-star or Oregon-Green (NEB) for 30 minutes 24 hours post plating. Media was removed and cells were washed with PBS. Cells were then treated with SNAP-Block (NEB) for 30 minutes. Cells were allowed to cycle for a subsequent 24 hours and then were treated with either DMSO or inhibitors, as noted. Cells were then fixed as noted above, or inhibitors were washed out and cells allowed to cycle and then fixed. Quantification of fluorescence intensity at centromeres was performed on single-stack images acquired at 60X or magnification and analyzed in FIJI (ImageJ). Centromeres were identified using the anti-centromere antibody (human anti-ACA, Antibodies.com) and a circle with approximate diameter of 0.2μm was used to measure mean SNAP intensity.

### Quantification of Fluorescence Intensity at Centromeres

Quantification of fluorescence intensity at centromeres was performed on single-stack images acquired at 60X magnification and analyzed in FIJI (ImageJ) using a custom macro. Interphase nuclei were segmented by thresholding the DAPI channel using the Huang algorithm, followed by shape smoothing. Nuclear ROIs were generated using the Analyze Particles function, and filled masks were used to define nuclear boundaries. Centromeric foci were identified by applying a bandpass filter to the centromere marker channel (ACA). Thresholding (Yen method) was applied to refine centromere foci. For each nucleus, background-subtracted intensity measurements were obtained from the measurement channel (e.g., Rad51) at centromere foci. ROIs lacking detectable centromeric signal above a minimum threshold were excluded from analysis. For each ROI, area, mean, and integrated fluorescence intensities were recorded.

Results were exported as CSV files and processed in R for downstream statistical analysis. All experimental conditions and biological replicates were batch-processed using identical segmentation thresholds and measurement parameters.

### Chromatin Immunoprecipitation

Cells grown in 150-mm dishes were treated with indicated drugs for 24 hours, released by treatment with 3 ml 0.05% Trypsin-EDTA (Gibco) 37°C for 10 min, neutralized by adding 6 ml DMEM-10% FBS, and centrifuged at 500x g for 5 min. Cells were resuspended in 6 ml ice-cold PBS (Gibco), cross-linked with 1% formaldehyde (16% formaldehyde, Electron Microscopy Sciences) for 10 min at r.t. The crosslinking reactions were terminated by addition of 125 mM glycine (Sigma), cells were pelleted by centrifugation at 600x g for 5 min, washed with 10 ml cold PBS. The cell pellets were incubated with 1 ml cell lysis buffer (20 mM Tris-Cl, ph8.0, 85 mM KCl, 0.5% NP-40, 1x protease inhibitors (Roche)) in ice for 10 min and centrifuged at 500x g for 5 min. The isolated nuclei were resuspended in 1 ml nuclei lysis buffer (50 mM Tris-Cl, ph8.0, 10 mM EDTA, 1% SDS, 1x protease inhibitors (Roche)) and sonicated to achieve ∼300- 500 bp fragments (Covaris) at 4°C for 24 min. The sonicated chromatin was centrifuged at 13000x g for 20 min at 4°C, aliquoted, diluted into 1 ml IP buffer (20 mM Tris-Cl, ph8.0, 150 mM NaCl, 1 mM EDTA, 1% Triton X-100, 0.01% SDS, 1x protease inhibitors (Roche)), and incubated with 2 mg primary antibody for 16h at 4°C. Protein G magnetic beads (Dynabeads, Invitrogen), 30 ml in 0.5% BSA-PBS (Gibco) were added to the reaction for 3h at 4°C. The beads were pelleted on magnetic block, washed sequentially with IP buffer, high salt buffer (20 mM Tris-Cl, ph8.0, 500 mM NaCl, 1 mM EDTA, 1% Triton X-100, 0.01% SDS, 1x protease inhibitors (Roche)), LiCl (20 mM Tris-Cl, ph8.0, 250mM LiCl, 1 mM EDTA, 1% NP-40, 1% Na deoxycholate, 1x protease inhibitors (Roche), and TE wash buffers, for 5 min at 4°C, 1 ml each wash, and eluted with 0.5 ml elution buffer (100 mM NaHCO3, 1%SDS) at 65°C for 20 min. The eluted DNA fragments were neutralized by addition of 40 mM TrisCl, ph 6.8, 1 mM EDTA, treated with 10 ug RNAseA (Invitrogen) for 30 min at 37°C, followed by incubation with 40 ug Proteinase K (Roche) for 60 min at 37°C, and purified using PCR purification kit (Qiagen). The purified DNA fragments were amplified in 20 ul qPCR reactions with iTaq Universal SYBR Green (BioRad) and primers corresponding to alpha-satellite regions of individual chromosomes using CFX96 Real Time PCR detection system (BioRad). The resulting Cq values of three biological replicates were averaged, normalized, converted to DNA concentrations and plotted relative to the input material.

### Subcellular protein fractionation

Cells were grown in 100-mm petri dishes until confluency, harvested by addition of 1 ml 0.05% Trypsin-EDTA (Gibco) and centrifuged at 500x g for 5 min. The pelleted cells were washed with 1 ml ice-cold PBS, and fractionated into cytoplasmic, membrane, nuclear, and chromatin fractions with Subcellular Protein Fractionation Kit (Thermo Scientific) according to manufacturer instructions. Briefly, cells were sequentially resuspended first in 100 μl cytoplasmic extraction buffer, incubated at 4°C for 10 min; then in 100 μl membrane extraction buffer, and centrifuged at 500xg for 5 min. The resulting cell pellets were incubated with 50 μl nuclear extraction buffer at 4°C for 30 min, centrifuged at 5000x g for 5 min and the supernatant (Nuclear fraction) was collected. Isolated nuclei were suspended in 50 μl chromatin extraction buffer in the presence of 10 mM CaCl2, 300 u MNase (Thermo Scientific), incubated at 37°C for 10 min, centrifuged at 16000xg for 5 min, and the supernatant (Chromatin fraction) was collected. Samples from Nuclear and Chromatin fractions were resolved of 12% SDS-PAGE gels, transferred to PVDF membranes (Millipore) and processed for Western blot analysis for the presence of indicated proteins as indicated.

### In vivo HU treatment

All described animal work was approved by the Institutional Animal Care and Use Committee (IACUC) of Yale University. CD1 mice were obtained from Charles River Laboratories. All mice were housed and maintained in a barrier facility with a 12 h light/dark cycle and ad libitum access to food and water.

In vivo HU treatment was performed as described previously^39^. HU was purchased from Sigma (H8627-5G) and resuspended in sterile water. Mice were intraperitoneally injected with 50 mg/kg body weight HU. After 6 hours, mice were euthanized and the jejunum was embed in Optimal Cutting Temperature (OCT, Sakura FInetek) and stored at -80°C. Frozen OCT blocks were sectioned at 8 µm thickness using a cryostat.

### Immunofluorescence of tissue

For cryosection staining, tissue sections were fixed for 8 min in 4% PFA, followed by a wash with PBS-T containing 0.2% Triton X-100 (AmericanBio). Sections were incubated in Mouse on Mouse Blocking reagent (Vector Laboratories) for 1 hr at RT. Primary antibodies were added to sections for overnight at 4°C. The primary antibodies used were as follows: Rabbit-anti CENP-A antibody (1:500 overnight, Invitrogen MA5-1429), Mouse-anti UBTF (1:1000 overnight, Abnova H00007343-M01) After washing in PBS-T secondary antibodies diluted 1:200 were applied to sections for 10 min. Secondary antibodies used were as follows: donkey anti-rabbit Alexa Fluor 488 (Jackson ImmunoResearch; 711-545-152), donkey anti-mouse RRX (Jackson ImmunoResearch, 715-295-151). DAPI (5 µg/ml; Invitrogen, D1306) was used as nuclear stain. After PBS-T washing, sections were mounted in antifade mounting medium (90% glycerol, 2.5 mg/ml *p*-Phenylenediamine in PBS).

Cryosections were imaged on an upright Zeiss AxioImager with Apotome 2 attachment and Zeiss AxioCam 506 mono camera using Zen software (v3.0; Zeiss). Objectives used were Plan Apochromat 10x/0.45 air, 20x/0.8 air, 40x/1.3 oil, and 63x/1.4 oil.

### Statistical Analysis and Significance Testing

In places where single comparisons were performed, we calculated significance (p-value) using a paired Student’s t-test. In places where multiple comparisons are performed, we used one-way ANOVA with Dunnett’s test when comparing multiple conditions to DMSO, or Tukey’s test when comparing across all conditions.

For all values, we normalized to the average value of the vehicle control for each biological replicate. No statistical method was used to predetermine the sample size. Data were excluded from analysis only in experimental replicates where positive controls failed to change in line with well-established expectations from prior literature. Experiments were not randomized and investigators were not blinded to experiments.

## Notes

### Competing Interest Statement

The authors have declared no competing interest.

