## Supplemental Figures for "ATR promotes genome instability via CENP-A eviction from centromeres under replication stress"

### Figure S1 Related to Figure 1

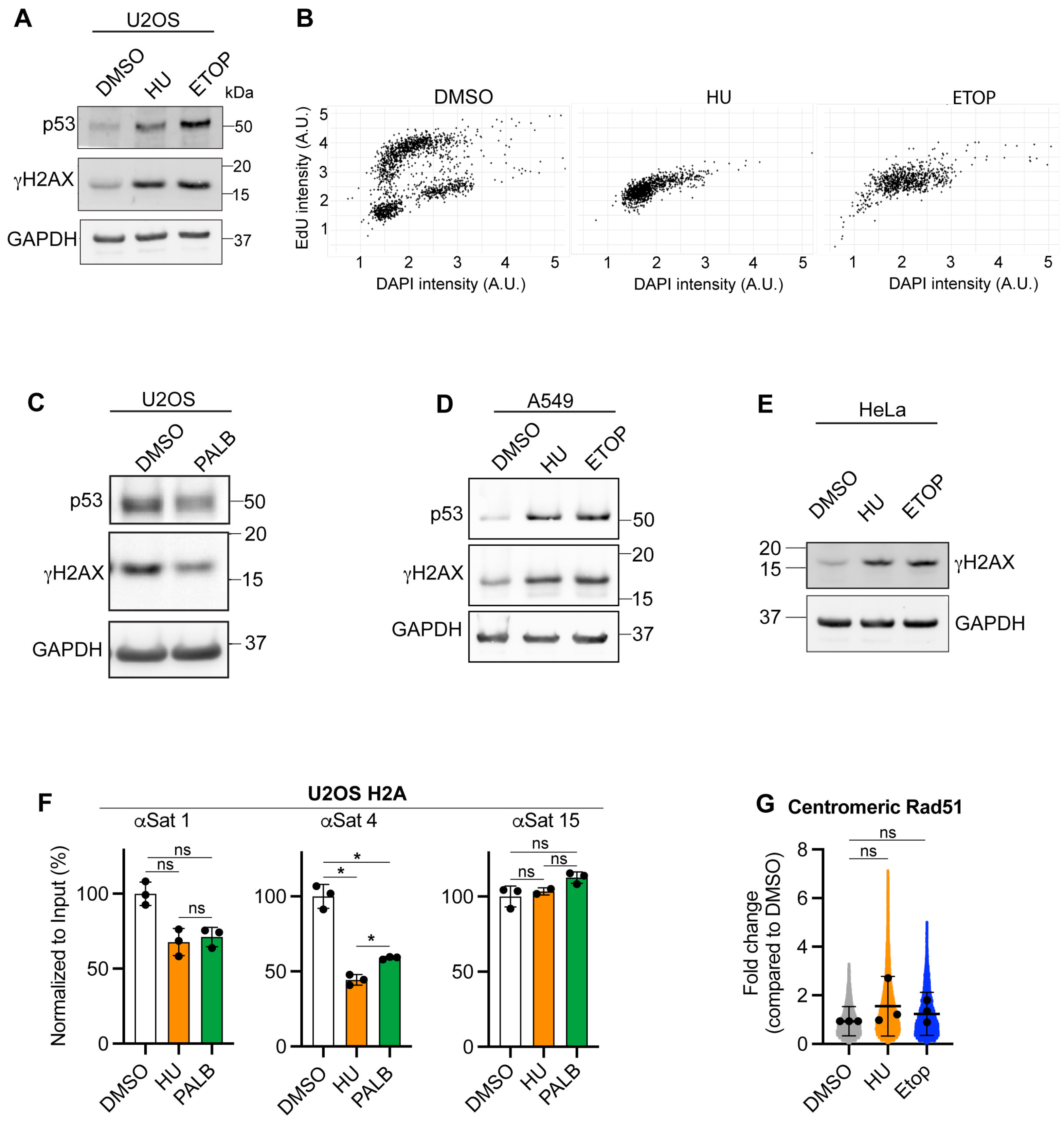

### Figure S2 Related to Figure 2

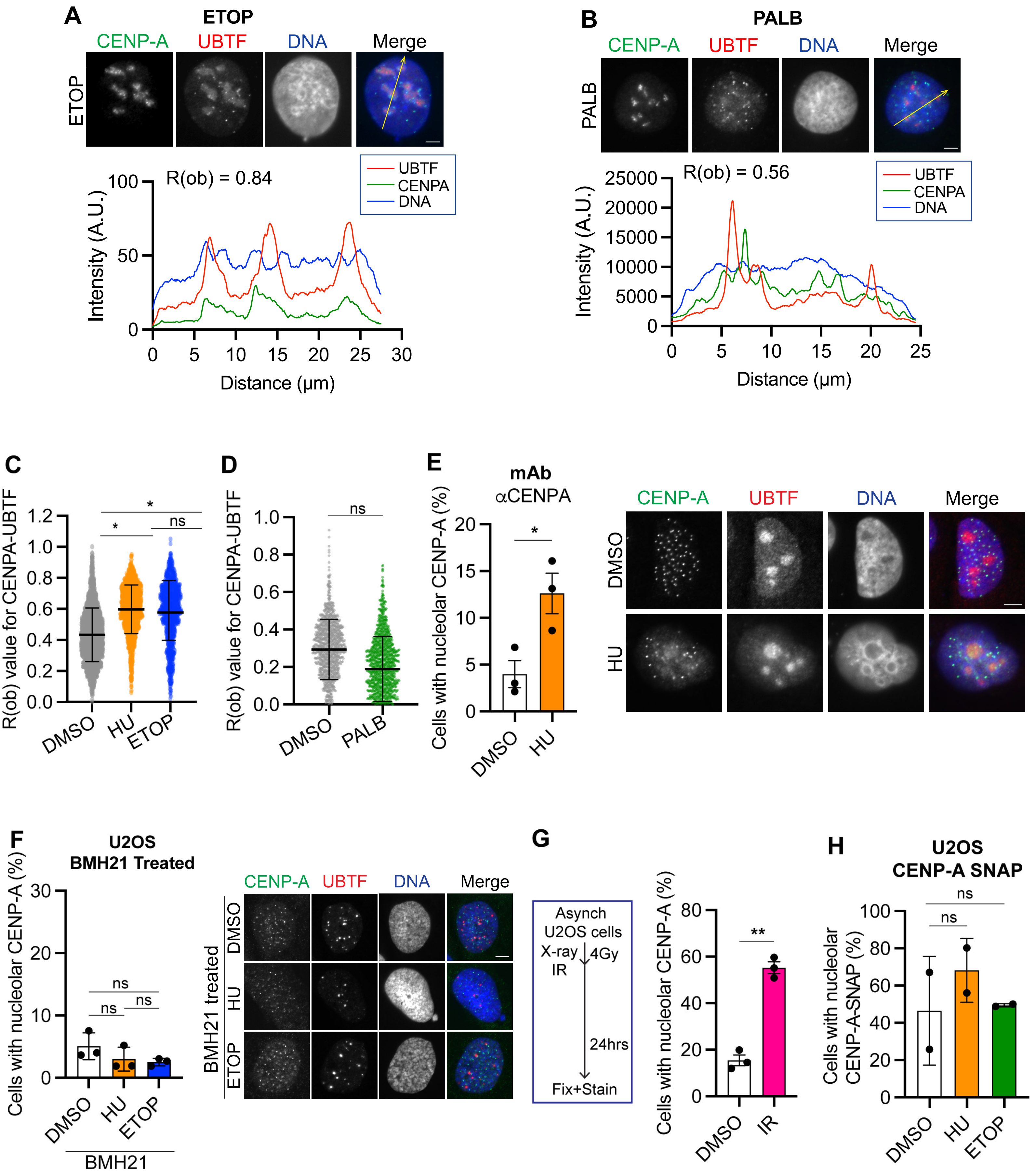

### Figure S3, related to Figure 3

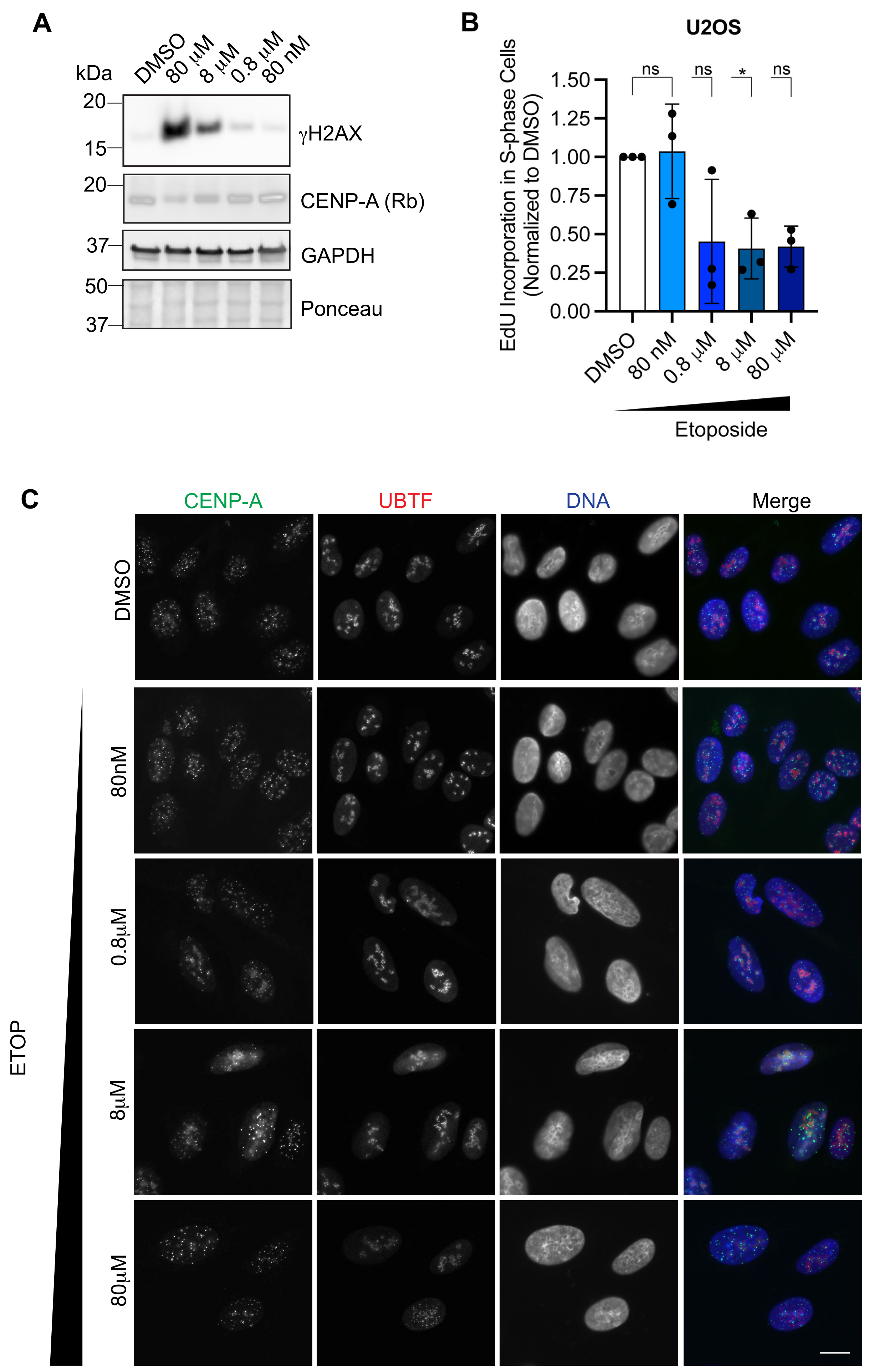

**Figure S4, related to Figure 3**

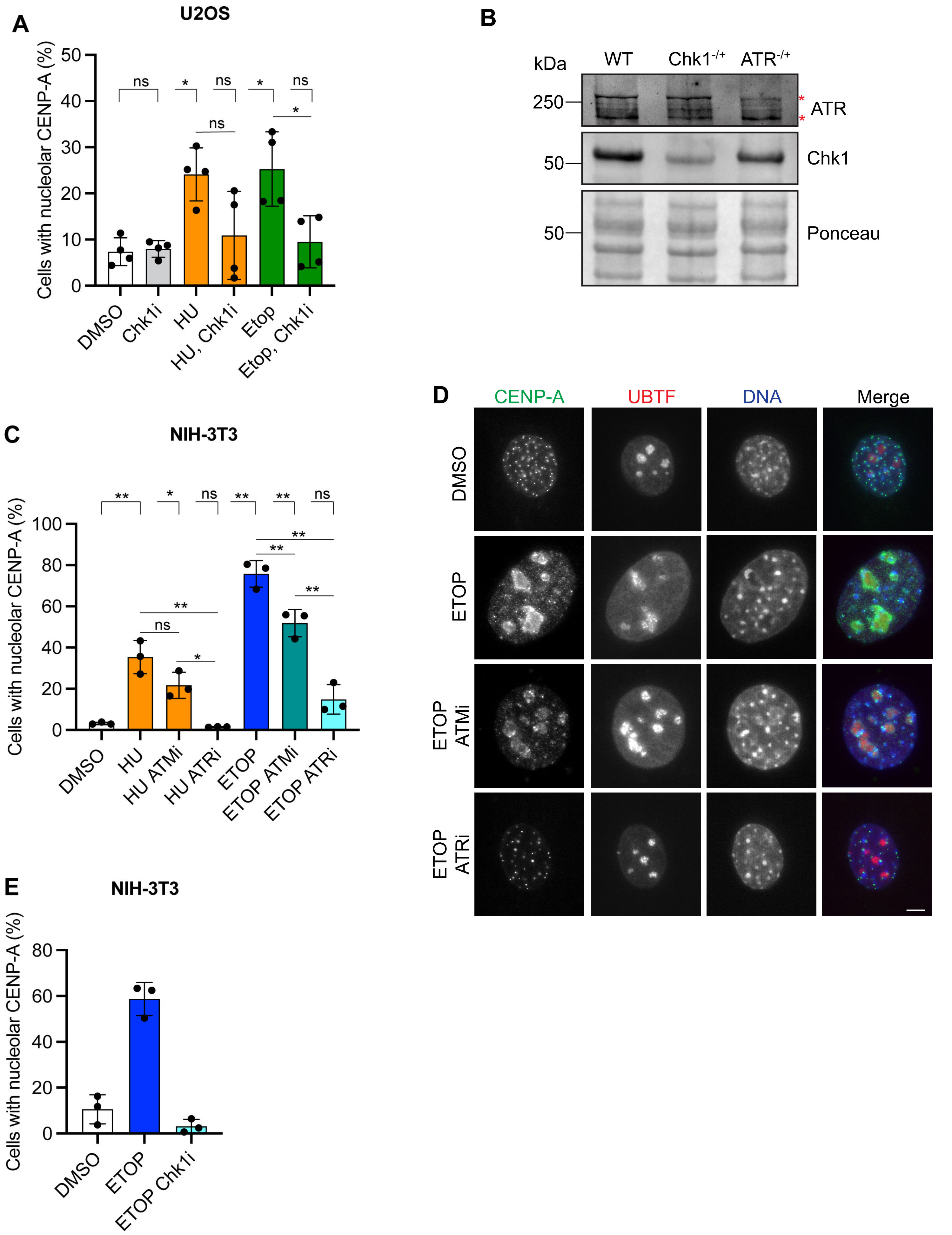

### Figure S5, related to Figure 4

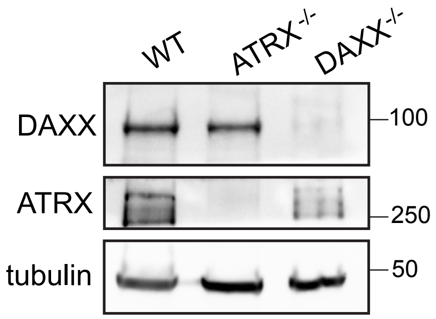

**Figure S6, related to Figure 5**

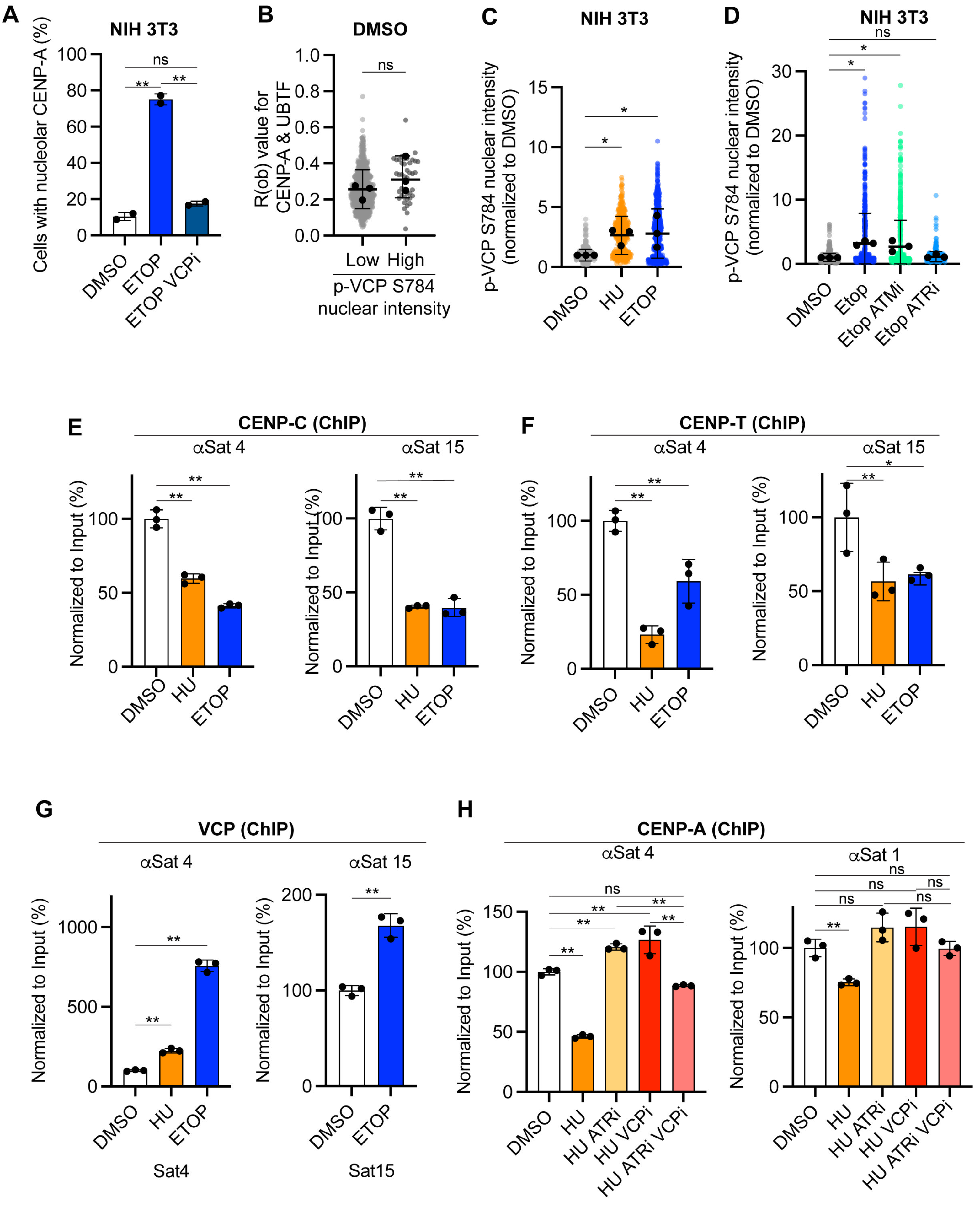
